# Structural basis of context-dependent RORγt target gene activation on chromatin

**DOI:** 10.1101/2025.03.10.642251

**Authors:** Timothy S. Strutzenberg, Xiandu Li, Matthew D. Mann, Hye Jeong Shin, Jordan Kelsey, Sriram Aiyer, Samuel M. Cotsarelis, Jingting Yu, Gennavieve Gray, Zeyuan Zhang, George Wanes, Zelin Shan, Bo Zhou, Ronald M. Evans, Paul C. Whitford, Patrick R. Griffin, Ye Zheng, Dmitry Lyumkis

## Abstract

RORγt is a key transcriptional regulator that supports protective pathogen clearance but also drives autoimmune disorders. Genetic studies have identified RORγt mutations that selectively protect from autoimmunity without compromising host defense. Herein, we describe the molecular principles governing how RORγt selectively drives these context-dependent gene programs. Structural studies revealed the selectivity-determining mutations lie in an α-helix, the C-terminal extension (CTE), which nucleates when RORγt engages partially occluded DNA response elements at nucleosome entry/exit sites. In *ex vivo* Th17 cells, the CTE is required for full genomic occupancy and expression of inflammatory effector genes, while residual DNA binding supports partial occupancy that, alongside context-dependent transcription factors, induces lineage-stabilizing gene expression. These findings support a model where RORγt uses distinct chromatin binding modes to engage an expansive chromatin landscape to activate target gene programs, explaining the complex phenotypes of CTE mutant mice.

## MAIN

Multicellular organisms rely on faithful regulation of gene expression to maintain homeostasis, whereas dysregulation of transcription contributes to disease. Signal transduction cascades activate transcription factors (TFs) that cooperatively engage non-coding DNA sequences in the genome called cis-regulatory elements (cREs) to control gene expression. The ENCODE consortium estimates that ∼2,000 TFs act through ∼2.4 million cREs to regulate ∼30,000 human genes^1^. A central challenge in gene regulation research is to understand how TFs elicit context-dependent functions in physiology and pathophysiology, which requires characterizing the genes that drive specific cellular phenotypes, identifying cooperating TFs and key cREs on which they operate, and defining molecular principles that govern gene expression activation.

The retinoic acid-related orphan receptor γ isoform 2 (RORγt) is a central regulator for different components of the adaptive immune system and plays many critical roles in physiology. RORγt is required for lymph node development^2,3^ and for thymocyte survival and T cell receptor recombination during thymopoiesis^3,4^. In peripheral T cells RORγt supports pathogen clearance by driving proinflammatory gene programs in IL-17-expressing T helper (Th17), γδ T cells, and Natural Killer T cells^5,6^. RORγt was recently found in antigen-presenting dendritic-like cells, where it is essential for antigen presentation and for tuning T cell responses^7^, implicating RORγt in tolerance as well as immunity.

RORγt also drives pathogenesis. Outside the immune system it supports survival and metastasis in triple-negative breast cancer^8^, pancreatic cancer^9^ and castration-resistant prostate cancer^10^, and its dysregulation drives autoimmune encephalomyelitis^11^, colitis^12^, type 1 diabetes^13^, and cancer progression^14^. These roles have driven development of RORγt antagonists for Th17-driven inflammatory and autoimmune disease^15^ and, more recently, agonists for antitumor immunity^16^. However, RORγt clinical trials have suffered high attrition from lack of efficacy or safety concerns^17^. A better understanding of cell-type-dependent RORγt function could therefore yield selective therapeutics targeting both autoimmune disorders and cancers.

Although RORγt is most studied in Th17 cells, cell-type specific RORγt functions are known. RORγt functions in Th17 cells by activating accessible enhancers that regulate expression of lineage defining genes such as *Il17a, Il1r1,* and *Il23r*. This is accomplished in concert with additional TFs, most notably BATF and IRF4, which remodel the chromatin landscape and mediate accessibility of RORγt to select genomic loci^18^. RORγt also interplays with RUNX1 and STAT3 to potentiate *Il17a* expression, both of which are required for complete activation^19,20^. Notably, a genetic screen identified an S92A/L93A double mutation that disrupts Th17 differentiation while preserving thymopoesis and some lymph node development^21^. How these processes operate at a molecular level remains largely unknown.

RORγt belongs to the nuclear receptor (NR) superfamily, which translates chemical signals into transcriptional changes. RORγt senses cholesterol metabolites^22,23^ and shares the modular domain organization of its NR relatives^24^. Its important elements include an N-terminal activation function-1 (AF-1) domain, a DNA-binding domain (DBD), a flexible hinge and a ligand-binding domain (LBD). The DBD recognizes ROR response elements (ROREs) in the conserved noncoding regions of the IL-17 enhancer^25^. The LBD contains the activation function-2 (AF-2) surface required for coregulator binding and ligand-dependent regulation^26^. RORγt activates transcription by working with steroid receptor coactivators (SRCs) and CBP/p300 acetyltransferases to engage chromatin and establish super-enhancers^27–29^. However, how the chromatin environment shapes RORγt binding, structure, dynamics, and regulation remains unclear.

Prior structural studies of RORγt examined individual domains. The sole RORγ DBD structure bound to a direct-repeat RORE defined DBD:RORE recognition and suggested dimerization on certain DNA sequences^30^, although the direct-repeat RORE used for structural studies contributes little to IL-17 expression, where RORγt acts as a monomer^25^. The LBD has been studied far more extensively, and structural work helped define both ligand- and coregulator-binding functions^31^ and explain recognition of hydroxycholesterol derivatives from the cholesterol and bile acid pathways^22,23,26^. While such efforts have shed light onto the activities of individual domains, they cannot capture how the domains act together in the full-length protein.

Here we investigate how the chromatin context shapes RORγt function. Using nucleosomes as the basic constituent of chromatin, we profile the sequence and structural determinants of RORγt-chromatin engagement and show by hydrogen-deuterium exchange mass spectrometry (HDX-MS) that nucleosome binding stabilizes the AF-2 coregulator surface while destabilizing the histone octamer. A cryogenic electron microscopy (cryo-EM) structure of the RORγt:nucleosome complex reveals a nucleated α-helix in the DBD C-terminal extension (CTE) and an opened entry-site DNA. Combining CTE mutagenesis in an *ex vivo* Th17 cellular model with interpretable base-pair-resolution analyses of genomic occupancy, we show that allosteric coupling to chromatin and crosstalk with neighboring TFs together govern context-dependent RORγt function. Collectively, we identify the RORγt CTE as a critical structural element that licenses RORγt to engage DNA that is partially occluded in chromatin to enact the RORγt target gene program.

## RESULTS

### RORγt preferentially engages nucleosomes at entry and exit sites

Like other NRs, RORγt is thought to bind preferentially at nucleosome-depleted genomic sites^18^. Yet RORγt must also bind chromatinized DNA to convert ‘closed’ to ‘open’ chromatin and establish the IL-17 enhancer^29^. Whether and how the chromatin environment influences RORγt binding remains unclear. To address this question, we profiled RORγt chromatin binding preferences using an assay called Selected Engagement of Nucleosomes coupled with sequencing (SeEN-seq)^32^, shown schematically in Fig. 1a. We tiled a classic RORE (cRORE; AAAGTGGGTCA) from the IL-17 enhancer^33^ and a variant RORE (vRORE; GTTCTGGGTCA) from the androgen receptor promoter^10^ through the tiling region of the W601 nucleosome positioning sequence. Assay optimization that reduced the detection of non-specific RORγt binding is described in Methods and Extended Data Fig. 1.

**Fig. 1:**
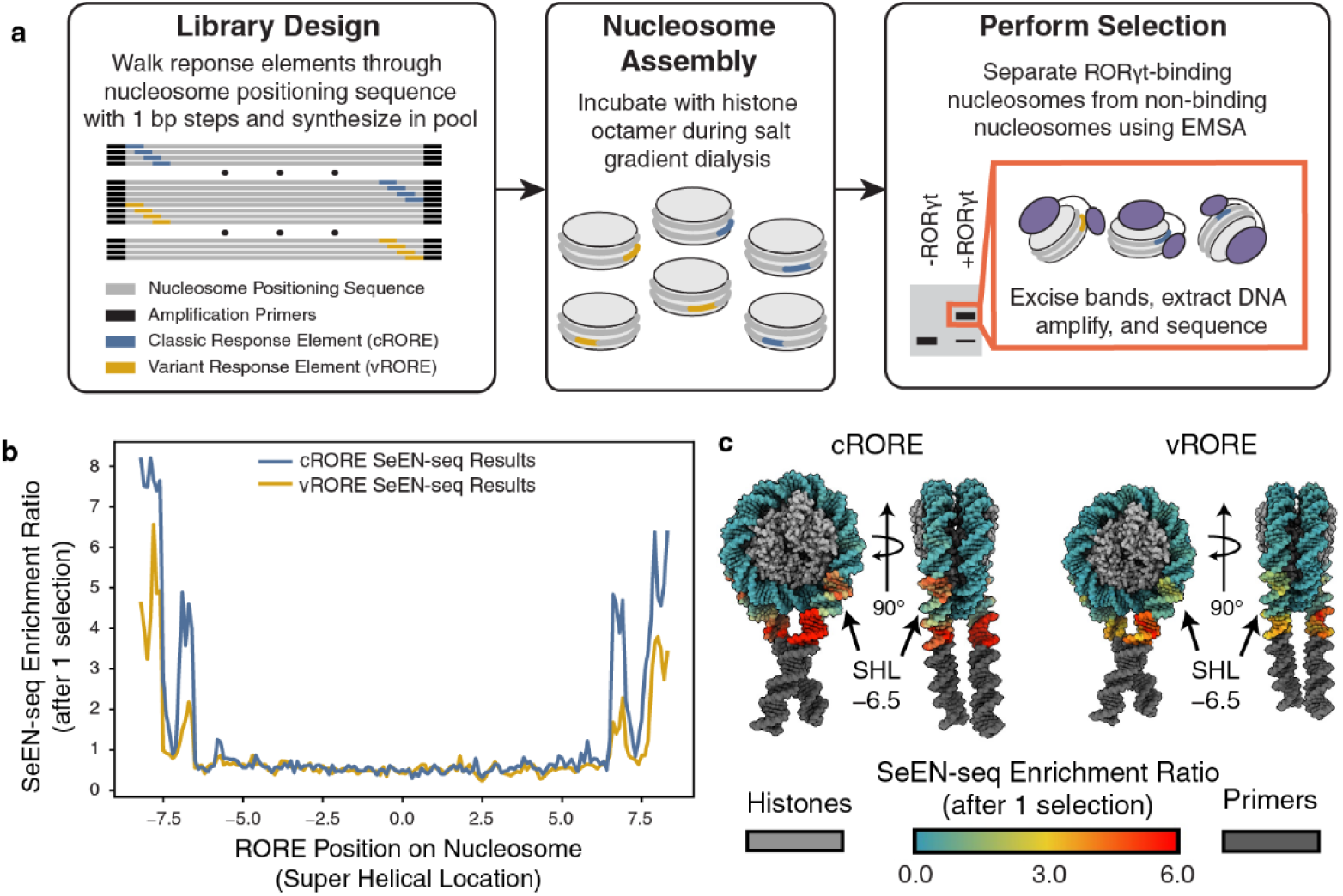
Sequence and chromatin context impact how RORγt engages response elements. (**a**) Schematic overview of the SeEN-seq assay. The cRORE and vRORE sequences are based on endogenous elements foud in the IL-17 enhancer^33^ and the Androgen receptor promoter^10^. (**b**) The results for tiling a cRORE or vRORE through the nucleosome positioning sequence after one round of selection. Log2 enrichment ratios are normalized to read counts from the library after selection divided by normalized read counts before selection. The RORE positions indicate where the center of the response element is located on the nucleosome in super helical location (SHL) notation. (**c**) The log2 enrichment ratios from SeEN-seq are painted onto a model of the nucleosome, generated using Alphafold3. The DNA bases are colored according to the color key legend at the bottom of the panel. Histones and primers are not covered and are colored accordingly.

The SeEN-seq experiment revealed the overall nucleosome binding preferences of RORγt (Fig. 1b). Beginning with the nucleosome entry site, binding is strongest at the DNA extension, inhibited at SHL −7.0, regained at productive sites centered on SHL ±6.5, and then progressively inhibited again toward the dyad. This pattern is inverted beyond the dyad, due to the pseudo-symmetry of the nucleosome. RORγt therefore prefers to bind ROREs at the extensions, entry, and exit sites of the nucleosome, with a periodicity of ∼10.5 bp. Access to internal ROREs between SHL ±5.5 is inhibited by the nucleosome core particle (NCP).

Mapping SeEN-seq signals onto the nucleosome showed enrichment where the major groove is exposed (Fig. 1c), consistent with known preference for major groove engagement^30^. When comparing RORγt-mediated cRORE versus vRORE engagement, we observe similar overall trends, but generally lower vRORE enrichment in the entry/exit sites, up to and including SHL ±6.5. This likely reflects lower affinity for vRORE, consistent with the reduced stability of RORγt bound to vROREs^34^. Thus RORγt binds entry/exit-site DNA with a periodicity that favors exposed major grooves.

### RORγt-nucleosome interactions promote changes in structural dynamics

For biophysical studies, we assembled a binary complex of full-length RORγt with a nucleosome carrying the cRORE at the entry site (SHL −6.5). The cRORE was chosen over the vRORE due to higher enrichment ratios by SeEN-seq, whereas the SHL –6.5 was chosen to maximize the likelihood of observing RORγt-nucleosome interactions, since cROREs beyond SHL ±7 are too dynamic, and those inside SHL ±5.5 are inaccessible. A 3:1 RORγt:nucleosome ratio yielded a stoichiometric complex that was purified by size-exclusion chromatography and confirmed intact by denaturing and native PAGE (Extended Data Fig. 2).

To determine how nucleosome engagement affects RORγt conformational dynamics, we used HDX-MS, which quantifies transient protein unfolding from backbone amide deuterium exchange^35^. We compared deuterium uptake in free versus nucleosome-bound RORγt across 32 peptides spanning 52% of the protein (Fig. 2a, Extended Data Table 1), and found structural changes throughout RORγt. As expected, the DBD core and CTE are protected (Fig. 2b-c), consistent with prior HDX-MS on free DNA^34^. Two observations were unexpected. First, the hinge between the DBD and LBD is deprotected, indicating increased dynamics upon nucleosome binding (Fig. 2d). Second, the LBD is protected across the conserved AF-2 coregulator surface at every time point measured (Fig. 2e). Prior HDX-MS characterization of RORγt conformational dynamics showed limited long range allostery between the DBD and the LBD upon DNA binding^34^, but those analyses used free DNA, which does not recapitulate a chromatin environment. At the same time, DBD-to-LBD allostery is documented in other NRs, including PPARγ:RXR^36,37^, RAR:RXR^38^, VDR:RXR^39–42^, LRH1^43^, and GR^44^. The observation that nucleosome binding via the RORγt DBD affects the distal AF-2 surface on the LBD suggests that RORγt is also regulated by long-range functional allostery, which is mediated by the chromatin context.

**Fig. 2:**
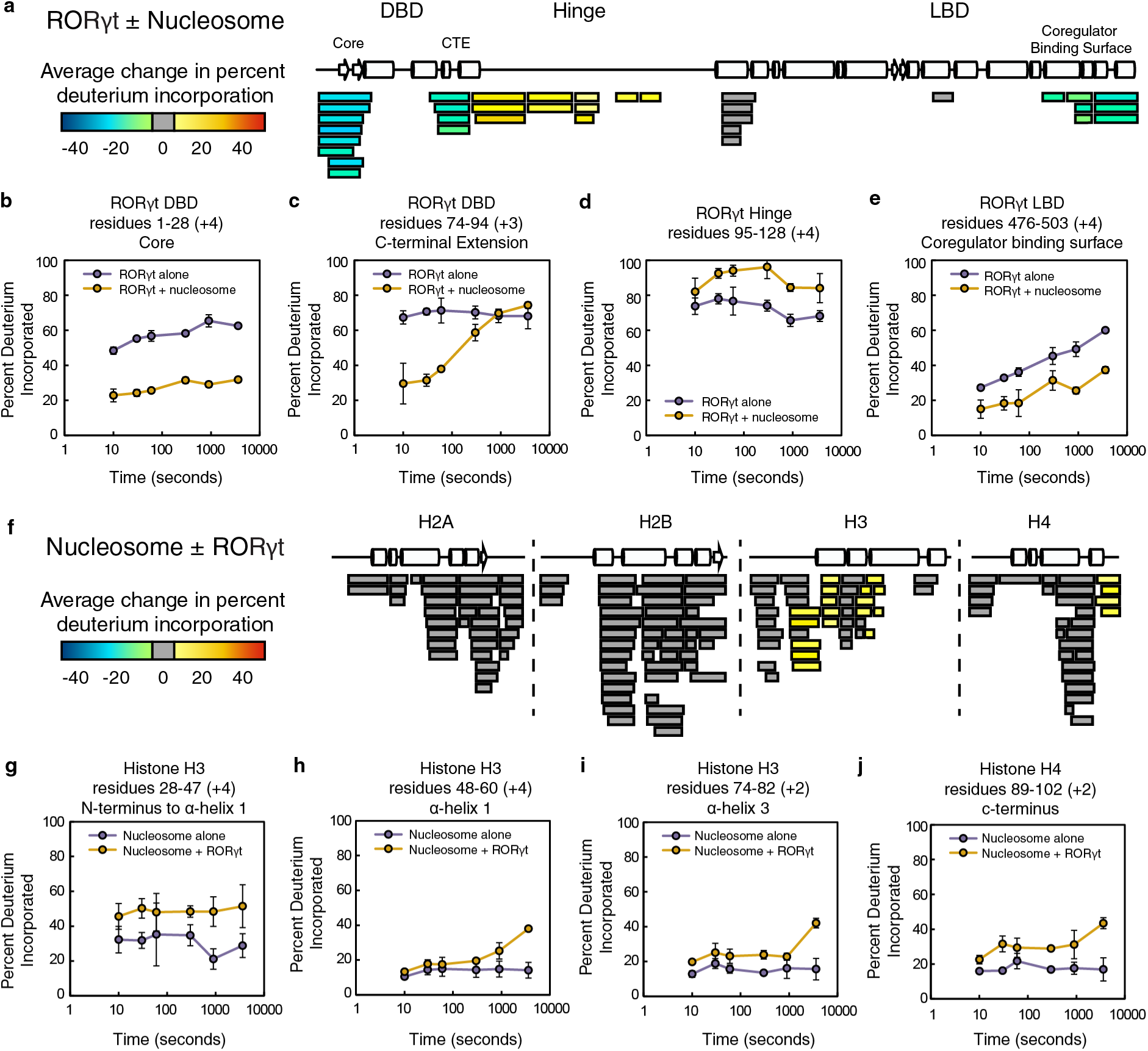
Interplay of structural dynamics between RORγt and the nucleosome. Results from HDX-MS analysis of RORγt ± nucleosomes are shown in panels **a**-**e**, while the results from HDX-MS analysis of nucleosomes ± RORγt are shown in panels **f-j**. (**a**) RORγt peptides that were quantified are shown as boxes below the primary structure of the protein. The color key indicates the average change in deuterium incorporation at all timepoints. Secondary structural elements are marked such that α-helices are cylinders and beta sheets are arrows. Selected deuterium build-up plots are shown for residues (**b**) 1–28 in the DBD, (**c**) 74–94 within the CTE, (**d**) 95–128 in the hinge, and (**e**) 476–503 within the AF-2 coregulator binding interface. The percentage of incorporated deuterium is plotted as a function of time. (**f**) Peptides for histone proteins were quantified and shown as boxes below the primary structures of each protein. Selected deuterium build-up plots are shown for residues (**g**) 28–47 in the N-terminus to first α-helix, (**h**) 48–60 in the first α-helix of H3, (**i**) 74–82 in the third α-helix of H3, and (**j**) 89–102 in the c-terminus of H4.

We next examined histone dynamics upon RORγt binding. Digestion optimization led to 85-95% coverage of all four histones (Fig. 2f, Extended Data Table 1). Deuterium uptake was unchanged for histones H2A and H2B, but there were localized deprotections to histones H3 and H4. We observed an increase in deuterium incorporation at the first and third α-helices of histone H3, Fig. 2g-i. There was also deprotection to histone H4 C-terminal tail, Fig. 2j, which contacts the H2A/H2B dimers within the NCP core. Most deprotections on histones H3 and H4 occurred at later timepoints, suggesting that RORγt binding dynamically destabilizes the histone octamer at these positions. As flexibility in the H4 C-terminal tail contributes to nucleosome remodeling^45^, deprotection of the H4 tail may serve as one of several cues that promote chromatin decompaction for transcriptional activation.

### Molecular architecture of the RORγt:nucleosome complex

To define the architecture of the binary complex and the interactions governing RORγt-nucleosome engagement, we determined its structure by cryo-EM (Extended Data Fig. 3; Extended Data Table 2), yielding a map resolved to resolutions ranging from 2.6 Å within the NCP to ∼6 Å in the entry-site DNA containing the cRORE and RORγt (Extended Data Fig. 4).

Density for the RORγt DBD is clearly defined within the nucleosome entry site that contains the cRORE (Fig. 3a). This enabled derivation of an atomic model for the DBD:cRORE interaction using integrative modeling with AlphaFold3^46^ and ISOLDE^47^ (Fig. 3b). There is additional density that resides above the histone octamer, but this density lacks discernable features and disappears at higher thresholds (Extended Data Fig. 4d–e). The limited resolution precludes *de novo* model building, but the size and shape of the density are consistent with the LBD. AlphaFold3 predictions likewise dock the LBD onto the histone octamer face while varying in its exact placement (Extended Data Fig. 5).

**Fig. 3:**
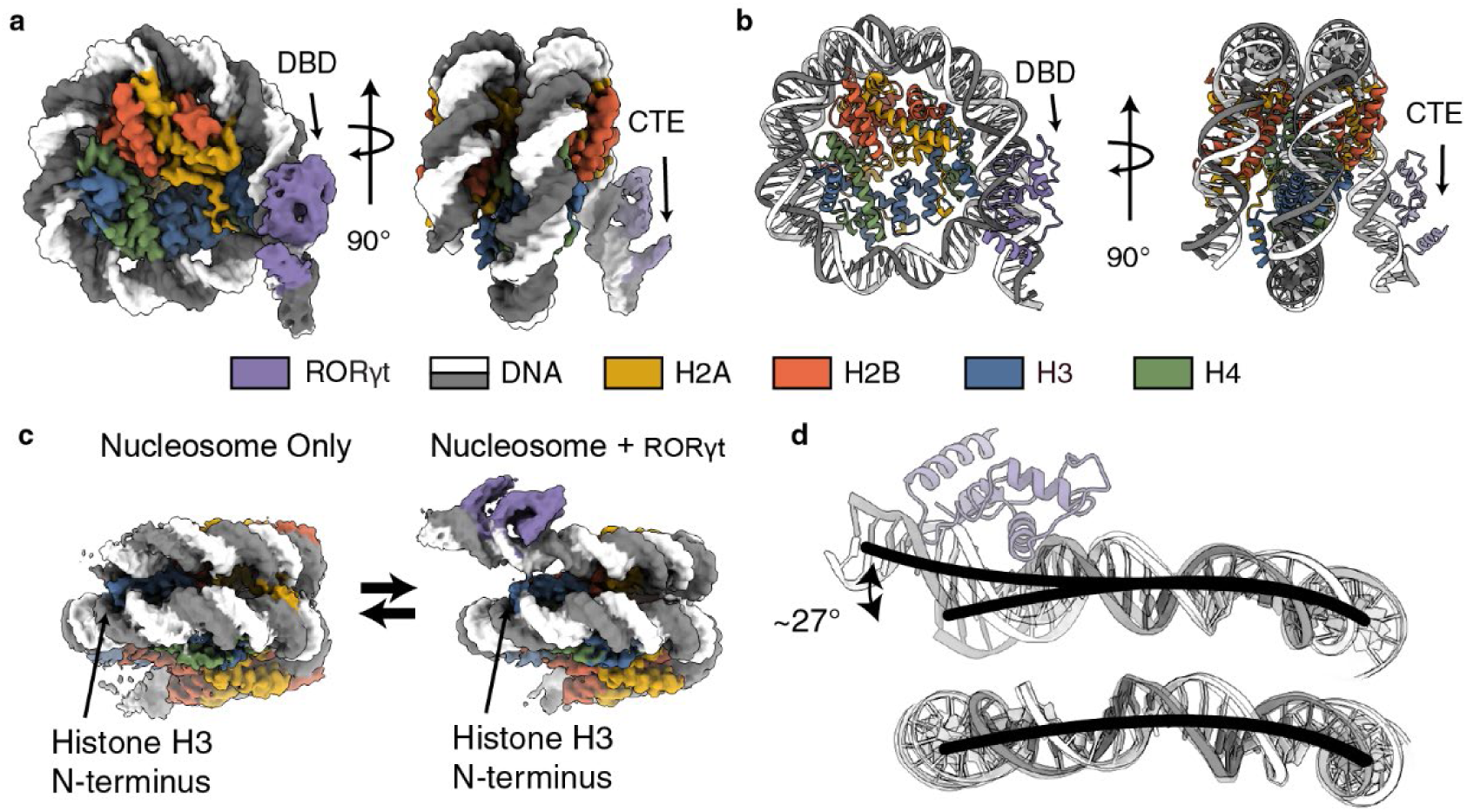
Molecular architecture of an RORγt:nucleosome complex. (**a**) The cryo-EM electron potential map of RORγt bound to the entry site of the nucleosome at SHL −6.5 is shown in two views. (**b**) Atomic models derived from potential maps are shown in the same views. (**c**) A comparison of a reconstruction of the nucleosome only and RORγt bound nucleosome. (**d**) atomic models for DNA and RORγt are compared. The DNA is traced with a black line to highlight differences.

The complex shows a reconfiguration of the entry-site DNA. To confirm that this results from RORγt binding, we determined a W601 nucleosome-alone structure at 3.1 Å for comparison. In the presence of RORγt, the entry site DNA is lifted off the histone H3 N-terminal tail and rotated by ∼27°, which is not observed in isolated W601 nucleosomes (Fig. 3c–d). This indicates that RORγt can induce and/or stabilize partial chromatin ‘opening’ when engaging ROREs at the entry site. Given the pseudo-symmetric engagement seen by SeEN-seq (Fig. 1b), the same reconfiguration presumably occurs at exit sites.

### Identification of a nucleated CTE within the RORγt DBD

RORγt uses its DBD to specifically bind the cRORE, which is located within the outward facing major groove at nucleosome SHL −6.5 (Fig. 4a). Helix 1 of the DBD inserts into the major groove, with structure- and sequence-specific contacts made by the first of its two zinc fingers. Lys31 confers sequence specificity by contacting the two guanine bases of the cRORE (arrow, NAANTAG↓GTCA; Fig. 4b). Outside the DBD core, RORγt makes minor groove contacts through its ‘KGFR’ motif, consistent with previous structural analyses^30,48^. In the context of the nucleosome, this motif inserts directly into the minor groove at SHL +7.0. Downstream, a prominent α-helix corresponding to the RORγt CTE stretches away from the nucleosome DNA.

**Fig. 4:**
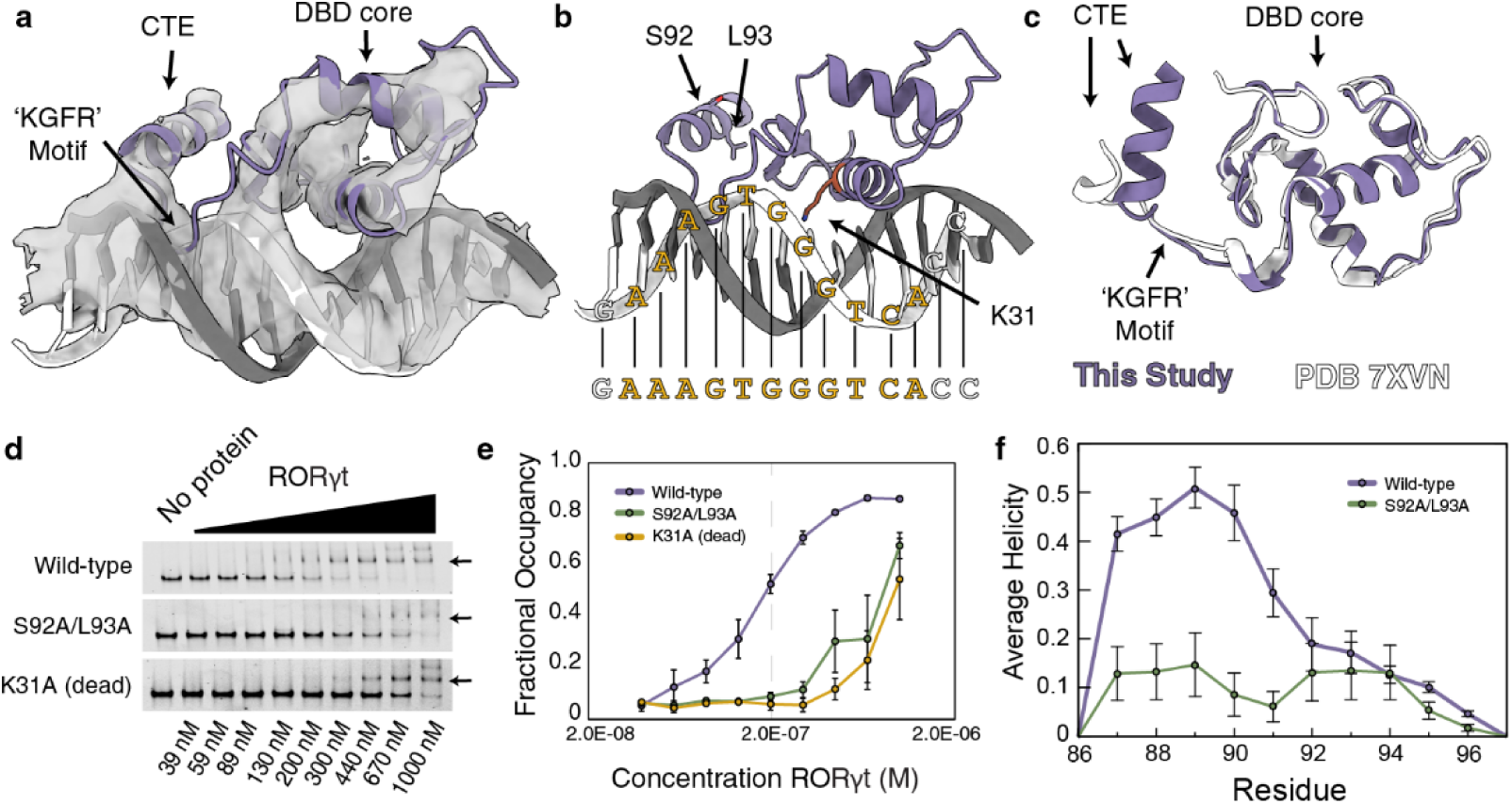
Identification of the RORγt C-terminal extension (CTE). (**a**) Close-up showing the experimental cryo-EM map and the atomic model of the RORγt DBD bound to the cRORE. (**b**) The model of the RORγt DBD containing residues S92 and L93 selected for mutagenesis and follow-up chromatin binding studies. Lysine 31 is displayed in red. (**c**) comparison of the DBD region from the cryo-EM model to a previously published crystal structure of RORγ (isoform 1) bound to a DR2 response element^30^. (**d**) Representative EMSA results for the binding of RORγt WT, and mutant proteins S92A/L93A or K31A to a nucleosome containing the cRORE are shown. The RORγt shifted bands are marked by black arrows. (**e**) The quantitation of 3 assay replicates are plotted with fractional occupancy on the y axis and protein concentration on the x axis. (**f**) Average per-residue helicity, calculated from explicit-solvent MD simulations for the WT (purple) and S92A/L93A (green) mutant. Error bars represent the standard error calculated from three independent 2-microsecond simulations.

In the structure of RORγ1 DBD (which has an additional 21 N-terminal residues compared to RORγt) bound to a DR2 element^30^, only the first four CTE residues are helical (Lys108–Arg111; Lys87–Arg90 in RORγt). By contrast, when RORγt is bound to the nucleosome containing a cRORE, residues spanning Lys87–Val97 within the CTE and the hinge region are clearly helical, Fig. 4c. AlphaFold3 models likewise predict a CTE α-helix (Extended Data Fig. 5), and HDX-MS shows strong protection across residues 74–94 upon DNA recognition (Fig. 2c), indicating that the CTE helix nucleates on cRORE binding.

The nucleation of the CTE α-helix has important implications for RORγt signaling. Previously, a genetic screen identified a double mutation S92A/L93A that distinguishes two separate biological functions mediated by RORγt^21^. In a model of experimental autoimmune encephalomyelitis (EAE), RORγt^S92A/L93A^ mutant mice were resistant to Th17-mediated autoimmunity but maintained largely normal thymocyte development and recovered lymph node biogenesis. Furthermore, these mice did not develop thymic lymphoma, which otherwise arises rapidly in RORγt^-/-^ mice^3^. Why these phenotypes diverge has remained unclear. Because our structure places S92 and L93 within the CTE, where they likely contact DNA, we hypothesized that the double mutation impairs RORγt chromatin binding and downstream activation.

We first tested the binding of WT, S92A/L93A, and K31A (DNA binding dead) mutants to nucleosomes containing cRORE at SHL −6.5 using an electrophoretic mobility shift assay. WT RORγt shifted nucleosomes with an EC_50_ of ∼100 nM, whereas the S92A/L93A and K31A mutants bound poorly and did not fully shift nucleosomes (EC_50_ > 1 μM; Fig. 4d-e). The CTE is therefore necessary for high-affinity chromatin binding in the absence of other factors.

To gain insights into the physical basis for the elevated EC_50_ values for the S92A/L93A mutant, we performed all-atom explicit-solvent molecular dynamics simulations of the wild type and mutant forms of RORγt bound to cRORE. For each system, we performed triplicate 2-microsecond simulations and calculated the average helicity within each residue of the CTE. These simulations indicate that the S92A/L93A mutation leads to a significant decrease in CTE helicity (Fig. 4f). In addition, its orientation is rotated relative to the WT protein, with the distribution of angles shifted by ∼40 degrees (Extended Data Fig. 6). This suggests that the S92A/L93A mutations energetically favor a more disordered CTE, explaining the loss of high-affinity chromatin interactions.

### The RORγt CTE contributes to Th17 differentiation but not IL-17 enhancer localization

We investigated the roles of the DBD core and CTE activities in a model Th17 system in which RORC-knockout CD4+ T cells are reconstituted by retroviral transduction with FLAG-tagged RORγt constructs and polarized with TGFβ and IL-6 (Fig. 5a). All constructs expressed normally, with the mutants showing ∼70% expression levels compared to the WT protein (Fig. 5b–c). WT RORγt induced IL-17A and completed Th17 differentiation, whereas K31A, S92A/L93A and empty vector (EV) negative control all showed greatly reduced IL-17A (Fig. 5d–e). These results support previously reported observations^21^, although the K31A mutant was not explicitly tested in the past. We therefore conclude that both the CTE and the DBD core DNA binding activities are necessary for activation of IL-17A.

**Fig. 5:**
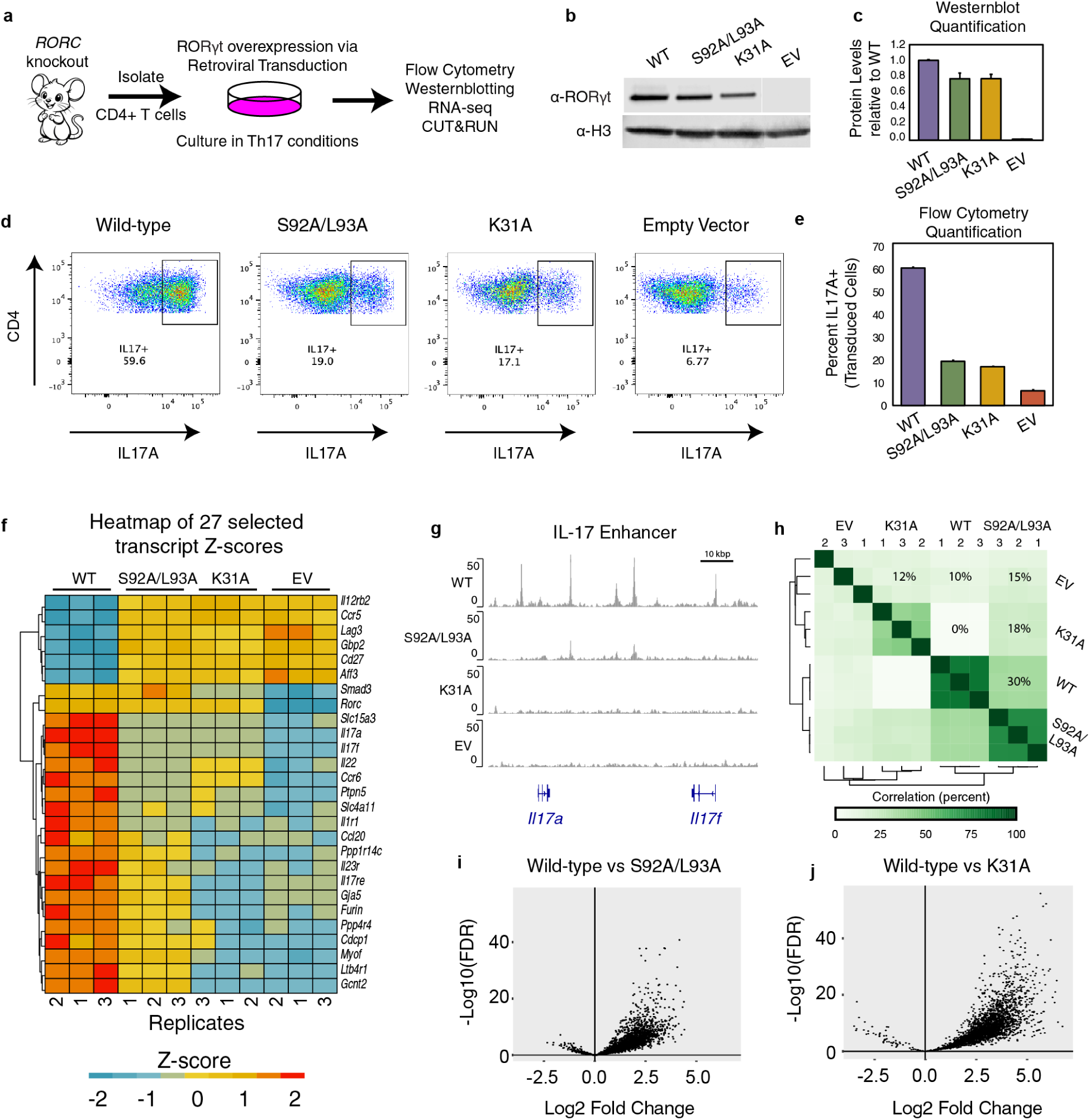
The RORγt CTE is required for inducing the Th17 gene program. (**a**) Schematic depicting our model Th17 cell culture system. (**b**) Representative Western blotting results and (**c**) the quantification of 3 biological replicates are shown. (**d**) Representative flow cytometry analysis of transduced CD4+ T cells and (**e**) the results of 3 replicates shown as a bar chart. (**f**) Results for RNA-sequencing is shown as a heatmap. A selection of 27 gene transcripts read counts are normalized to Z scores. (**g**) CUT&RUN was performed for 3 biological triplicates, the alignment results were merged, and read peaks are shown at the IL-17 locus. (**h**) Quantificaiton of peak intensity from CUT&RUN data using DiffBind analysis, plotted as a correlogram. Differential analysis comparing (**i**) WT with S92A/L93A, and (**j**) WT with K31A, both shows as volcano plots, respectively.

To assess the impacts of RORγt mutations on the Th17 differentiation program, we used transcriptomic analysis by bulk RNA sequencing (RNA-seq). Comparing the WT and EV controls, we found that WT RORγt overexpression leads to significantly increased transcription of 222 genes – including key Th17 lineage defining genes *Il17a*, *Il17f*, *Il23r*, *Ccr6*, and *Cxcl3* – and decreased transcription of 218 genes (Extended Data Fig. 7a). We selected 27 genes with the most significant differential RNA expression that also exhibit RORγt genomic occupancy near the gene body (Fig. 5f). These include classic targets (*Il17a*, *Il23r*, *Ccr6*), genes newly induced by RORγt overexpression (*Furin*, *Cdcp1*, *Gja5*) and genes that are downregulated (*Il12rb2*, *Ccr5*, *Lag3*), which generally lack RORγt occupancy at the gene body. Core DBD activity was necessary for both occupancy and activation. We also find that the S92A/L93A RORγt mutant weakly transactivates a subset of target genes like *Il17re, Il23r, and Furin*, unlike the DNA-binding dead RORγt and EV control, although transactivation was attenuated relative to WT (Extended Data Fig. 7b-d).

We next assessed genomic occupancy by Cleavage Under Targets and Release Using Nuclease (CUT&RUN). WT RORγt bound the IL-17 locus in line with published datasets (Extended Data Fig. 8a), whereas K31A showed no binding (Fig. 5g). The S92A/L93A mutant shows similar trends in genomic occupancy as the WT protein, albeit with reduced signal (Fig. 5g). Differential binding analysis confirmed that S92A/L93A correlates poorly with EV and K31A but well with WT (Fig. 5h). Overall, the CTE mutations reduced RORγt genomic occupancy (Fig. 5i) but less severely than K31A (Fig. 5j) and did not substantially change where RORγt binds. These results indicate that S92/L93 are required for high-level *Il17a* expression but not for occupancy at targets such as *Il23r*, *Furin*, *Gcnt2* and *Ltb4r1*. The CTE therefore regulates genomic occupancy in a context-dependent manner.

### Sequence determinants for RORγt genomic occupancy at cis regulatory elements

How can reduced *in vitro* chromatin affinity be reconciled with widespread genomic occupancy in cells? S92A/L93A RORγt binds chromatin poorly *in vitro* and is modestly less abundant, possibly reflecting increased ubiquitin/proteasome-mediated degradation^21^, yet it still engages thousands of genomic loci, with context-dependent effects on activation.

To gain deeper insight into the broader trends of our CUT&RUN datasets, we used a deep learning model called base pair network (BPNet) to decode TF binding motifs associated with RORγt, with single base-pair resolution. BPNet quantifies how sequence contributes to TF occupancy^49^. Models trained on WT and S92A/L93A CUT&RUN peaks (Fig. 6a) were used for motif discovery. The S92A/L93A mutant RORγt has reduced preference for the A-rich 5′ end of the motif compared to WT (Fig. 6b). Because the CTE extends from this end of the motif and is expected to be disordered in the mutant, this loss of preference makes sense. Further, the sequence preference of WT RORγt is more closely aligned with cROREs, whereas the sequence preference of S92A/L93A RORγt is more closely aligned with vROREs.

**Fig. 6:**
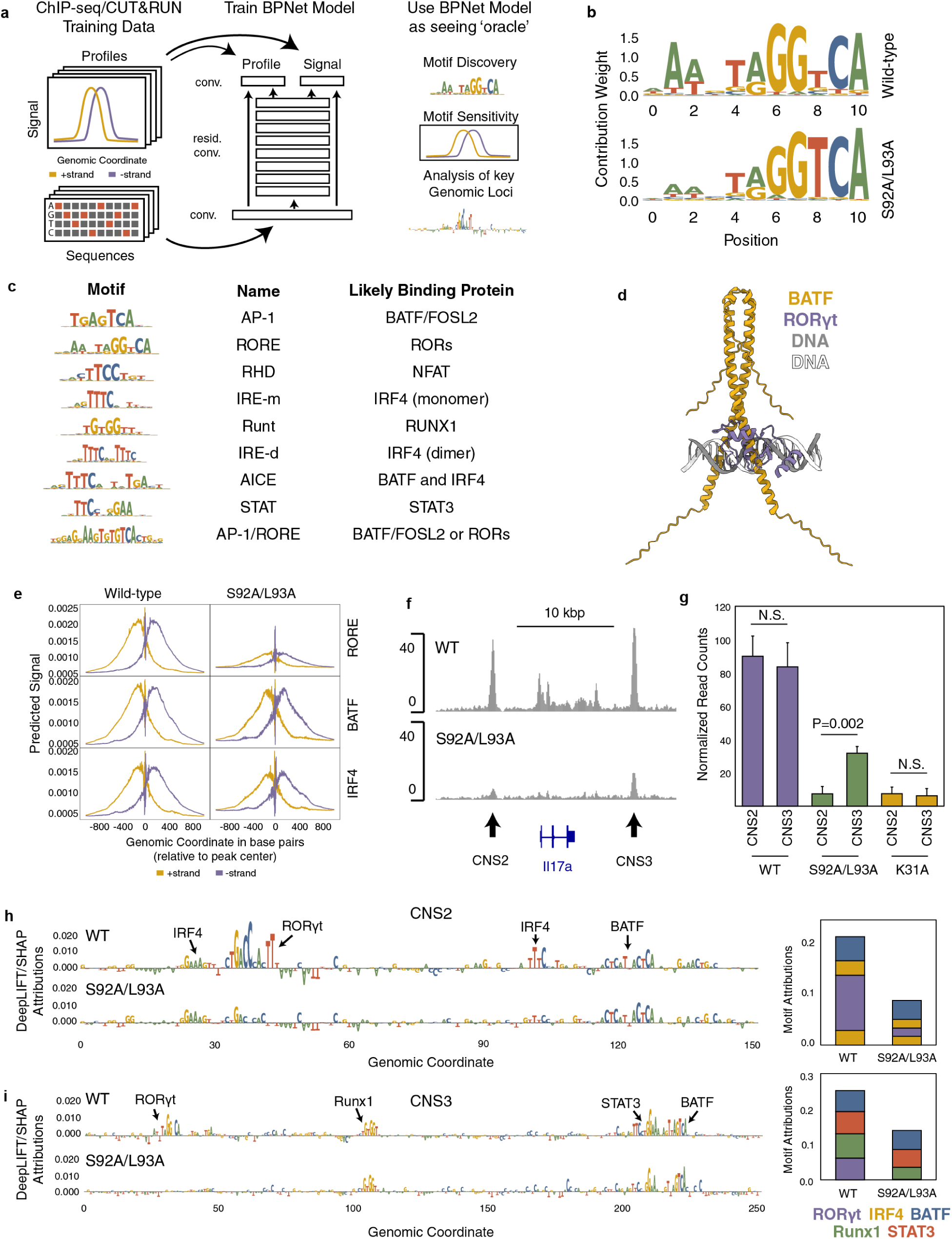
Sequence determinants for RORγt genomic occupancy in Th17 cells. (**a**) schematic showing the workflow for using BPNet to interpret genomics datasets. (**b**) We performed motif analysis using TF-modisco, and the results for the wild-type and mutant datasets are shown. (**c**) TF-modisco analysis of previously published ChIP-seq datasets revealed nine motifs of interest. (**d**) A novel motif was used for structure prediction with the RORγt DNA binding domain and a BATF dimer. The resulting structure predictions are aligned to the DNA sequence and suggest a competition for the DNA major groove between BATF and RORγt. (**e**) The results of marginalization experiments are shown. Briefly, GC-matched negative control sequences were mutagenized *in silico* to introduce a motif, and the change in predicted signal is plotted as a function of location in the sequence. The results shown are for 3 selected motifs (a cRORE, the AP-1/BATF motif, and the IRF4 dimer motif) and scored using models trained on WT or S92A/L93A datasets. (**f**) A zoom-in on ChIP-seq alignment tracks are shown at the IL-17a gene. Conserved noncoding sequnces (CNS) of interest are labeled. (**g**) Changes in peak intensity for CNS2 and CNS3 across our CUT&RUN datasets were quantified with DiffBind and shown as a bar plot. Attribution analysis was performed at (**h**) CNS2 and (**i**) CNS3. Briefly, genomic sequences were extracted and individual bases were mutagenized *in silico* and changes in score were determined using BPNet models trained on WT or S92A/L93A ChIP-seq datasets using DeepLIFT/SHAP. The change in score are plotted in the y axis as DeepLIFT/SHAP attributions as a function of genomic coordinate. The interpretation of the likely binding proteins are labeled above the motifs and indicated with black arrows. Inlaid plots on the right sum the attribution values of the corresponding motifs according to the BPNet models trained on WT and S92A/L93A ChIP-seq datasets.

We next asked how TF repertoires at different loci influence RORγt occupancy, and why some loci are more sensitive to CTE mutation. RORγt acts on cis-regulatory elements (cREs) cooperatively with other TFs, including but not limited to BATF/FOSL2, IRF4, STAT3, Runx1, and NFAT^18^. To capture these signals, we performed motif discovery using BPNet models trained on previously published ChIP-seq experiments^21^. These datasets are broader and deeper than our CUT&RUN data (Extended Data Fig. 8b; Extended Data Table 3). Motif discovery recovered many known RORγt collaborating motifs at cREs (Fig. 6c). We also found a mutually exclusive AP-1/RORE motif that can accommodate either bZIP proteins such as BATF/FOSL2 or RORγt; AlphaFold3 predicts that the two compete directly for the major groove at these sites (Fig. 6d).

To determine how individual motifs contribute to occupancy, we marginalized motifs into GC-matched non-peak background sequences and scored the predicted increase in signal for WT and S92A/L93A models. For BPNet models trained on the WT dataset, this analysis predicts strong signal for the RORE alongside motifs for IRF4 and BATF (Fig. 6e), consistent with the well-known cooperativity between RORγt and BATF/IRF4 at cREs^18^. However, for BPNet models trained on the S92A/L93A dataset, we see retention of sensitivity for IRF4 and BATF but a marked reduction in predicted signal for the RORE. This suggests that S92A/L93A RORγt is retained at ROREs by neighboring IRF4 and BATF, consistent with its reduced intrinsic RORE affinity *in vitro* and residual binding through the DBD core.

Finally, we examined individual cREs within the IL-17 locus. RORγt engages conserved non-coding sequences (CNSs) that act as cREs but contribute unequally to IL-17 expression; we focus on CNS2 versus CNS3 (Fig. 6f). CNS2 is functionally relevant, because its deletion reduces IL-17 expression and protects mice from Th17-mediated autoimmunity^25^. CNS2 also becomes hyperacetylated during Th17 differentiation and is located approximately 3 kbp (i.e. ∼20 linearly assembled nucleosomes) away from the IL-17a promoter. Less is known about CNS3, but it is interesting because the S92A/L93A mutant RORγt was somewhat retained at this site (Fig. 6f). Genomic occupancy is nearly ablated at CNS2 for the S92A/L93A mutant, indicating that the CTE is critical for binding to this location, but occupancy is less reduced at CNS3 (Fig. 6g). To investigate these differences, we performed attribution analysis using DeepLIFT/SHAP analysis^50,51^. We find that RORγt occupancy at CNS2 is driven primarily by a cRORE with little contributions from other TFs such as BATF and IRF4, despite the presence of their motifs (Fig. 6h). In the case of CNS3, we observe a greater contribution from other TF motifs including STAT3, Runx1, and BATF, as evidenced by higher attribution values relative to RORγt (Fig. 6i). Overall, these trends indicate that CNS2 likely requires high affinity RORγt/RORE interactions driven by the nucleated CTE to facilitate genomic occupancy and establish the IL-17 enhancer. By contrast, accessory TFs are predicted to play larger roles in promoting RORγt occupancy at sites like CNS3 to facilitate lower affinity interactions of RORγt with vROREs to maintain sufficient genomic occupancy.

## DISCUSSION

Our data show that both the DNA sequence bound by RORγt and nearby TFs that likely stabilize its binding on ROREs are determinants of chromatin occupancy and target gene expression. The results point to a complex network of allostery and highlight how the chromatin environment can influence RORγt structure, dynamics, and function. Based on the collective observations, we propose that the chromatin environment governs both RORγt CTE nucleation and AF2 stabilization (Fig. 7a). These activities may jointly promote coregulator recruitment, increase histone accessibility for post-translational modification by destabilizing nucleosome DNA, and facilitate enhancer formation.

**Fig. 7:**
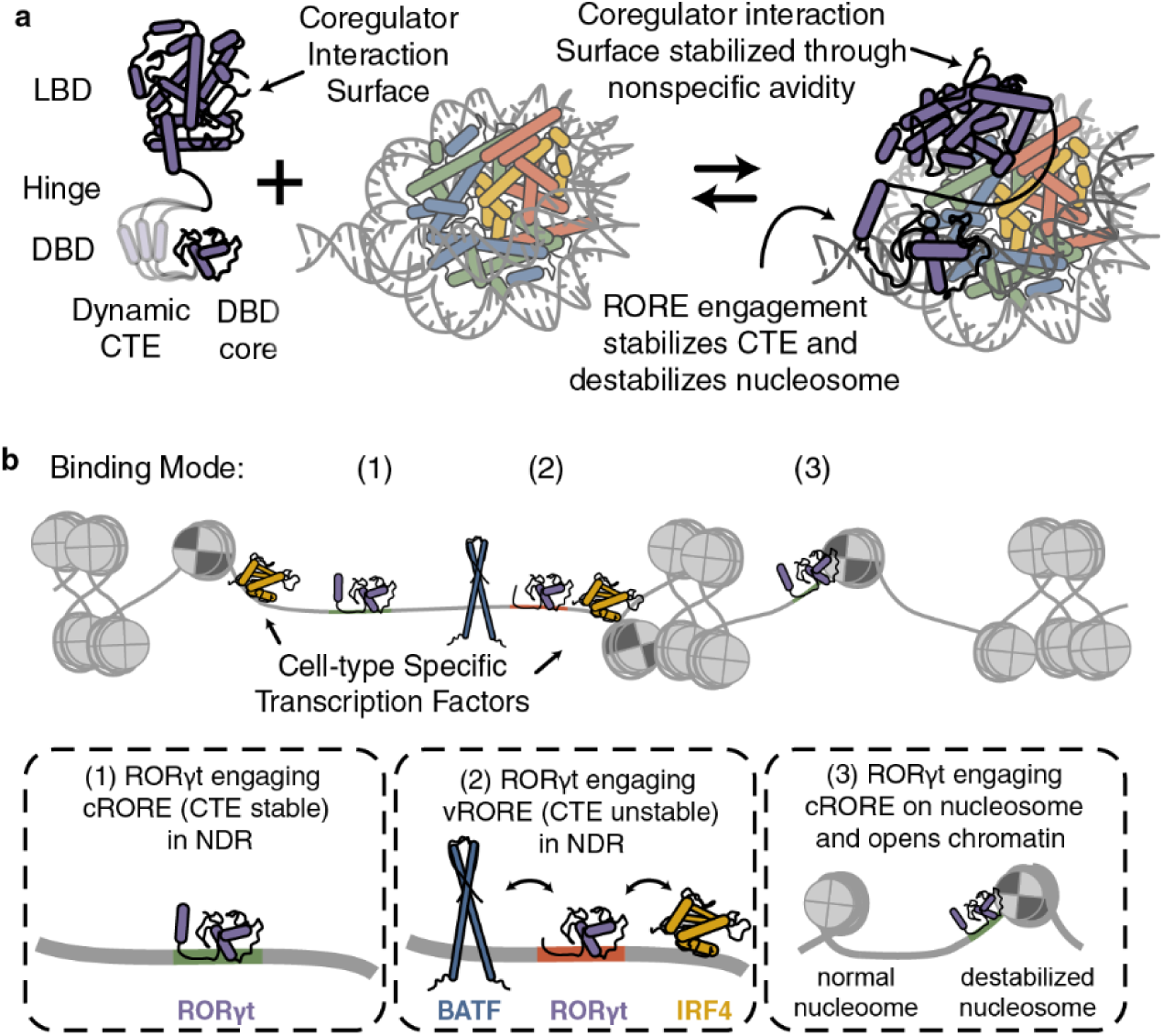
Model for RORγt genomic occupancy. (**a**) An updated RORγt activation model where nucleosome engagement simultaneously stabilizes the RORγt CTE and AF-2 coregulator binding surface while also destabilizing the nucleosome. (**b**) Cartoon schematic depicting different scenarios of RORγt engaging chromatin in the genome. In Th17 cells, the primary collaborating TFs are BATF and IRF4, which are shown in the schematic, whereas other TFs likely interplay with RORγt in different cell types. NDR indicates nucleosome depleted regions.

Genomic ROREs are the native targets for RORγt, yet the role of chromatin in shaping RORγt structure and function has been largely unexamined. Using SeEN-seq, we find that RORγt binds preferentially at nucleosome entry/exit sites and engages cROREs (AAAGTGGGTCA) with higher affinity than vROREs (GTTCTGGGTCA). The differences between these motifs are largely constrained to the 5′ motif sequence, which is A-rich in the cRORE and interfaces with the CTE. Mutation of the CTE relaxes RORγt specificity for cROREs, such that the recognition motif becomes more similar to vROREs. Consequently, CTE mutations cause an uneven loss of chromatin binding in Th17 cells, wherein some sites are more heavily impacted than others. Although chromatin states in Th17 cells are not mapped, these data suggest that sites most affected by CTE mutation (such as CNS2) carry less accessible ROREs, embedded near nucleosome entry/exit sites, whereas less affected sites (such as CNS3) carry ROREs in more accessible free DNA. The CTE is thus a key structural determinant of RORγt genomic occupancy.

The CTEs of monomeric NR DBDs sharpen sequence specificity, but their consequences in chromatin remain underappreciated. For example, the ‘RGGR’ motif located in the CTE in NR5A2 (LRH-1) was recently suggested to play a role in pioneering functions, but not in DNA binding activities^52^. For RORγt, CTE dynamics change with the bound DNA sequence^34^. Unexpectedly, our cryo-EM structure, supported by HDX-MS and by all AlphaFold predictions, reveals an α-helical CTE that is likely nucleated by the ‘KGFR’ motif engaging the 5′ A-rich extension present in cROREs but not vROREs. A vRORE-bound CTE would presumably be less helical, further shifting preference toward free DNA.

Like other NRs, RORγt must integrate many signals from the DNA sequence^34^, its post-translational modification status^53^, cooperating TFs on cREs^18^, and the nature and presence of bound ligands^22,23^, in order to recruit transcriptional coregulators and influence target gene expression. A hallmark of active RORγt is a stabilized AF2 surface, which small molecules can reinforce to increase coregulator affinity and ligand-dependent activity^26,54^. However, prior attempts to identify cross-domain allostery on free DNA in RORγt were unsuccessful^34^. Our HDX-MS observation that a nucleosome-embedded cRORE stabilizes this surface is therefore notable. Although the LBD density above the histone octamer is weak, precluding high-resolution insight into the AF2 conformation, the long 147-residue unstructured linker connecting the DBD to the LBD can conceivably promote transient LBD engagements with other nucleosomes and/or associated proteins in a more native chromatin environment or in the context of chromatin condensates. Long-range allostery between the DBD and the LBD is a well-known property of NR activation^44^, which we propose is also applicable to RORγt.

We propose a model for context-dependent RORγt function (Fig. 7b). In this model, ROREs are occluded by chromatin until some cell-type specific TF repertoire activates cREs and promotes the exposure of ROREs. RORγt then engages these ROREs with different affinities depending on their sequence and the chromatin context within which the RORE is embedded.

The three scenarios implied through our work include (1) RORγt engaging a cRORE alone within a nucleosome depleted region, (2) RORγt engaging a vRORE in concert with other TFs that stabilize the protein on DNA, and (3) RORγt engaging a cRORE near nucleosome entry/exit sites, which in turn partially destabilizes the nucleosome and exposes the histone octamer. Subsequently, coregulatory proteins interface with RORγt and other TFs to promote acetylation and impact transcription on adjacent target genes.

Can this model extend to other NRs? RORγt acts as a monomer that activates rather than represses transcription and, until now (Fig. 2), had not been shown to exhibit cross-domain DBD-LBD allostery. By contrast, the glucocorticoid receptor (GR), which is a more complex NR, adopts multiple oligomeric states, can activate or repress, and shows well-known DBD-LBD allostery^44^. To achieve high affinity DNA interactions, RORγt (like other monomeric NRs) relies on the CTE. GR also uses its CTE^55^, but can additionally tune high affinity DNA interactions through cooperative homodimerization on GR response elements (GREs)^56^. In both receptors, mutations that disrupt high-affinity DNA binding give intermediate phenotypes between null and WT. GR depletion is embryonic lethal, yet mice bearing GR mutations that disrupt only DNA-induced dimerization are viable^57^ and redistribute GR away from palindromic but not monomeric GREs^58^. We therefore posit that high-affinity DNA binding is required where response elements are less accessible but dispensable where they are not. Therefore, functional selectivity could be a product of chromatin sequence- and structure-dependent cues unique to different cellular environments.

RORγt has evolved to play diverse and complex roles in immunity. Its CTE is shared by all mammalian ROR family members and retains high sequence similarity with nematode HR3 receptors, placing its origin in the last common ancestor with bilaterians, which pre-dates adaptive immunity by ∼100 million years^59,60^. The most likely scenario is that its chromatin-reading dependence was already in place in ancient receptors, and that this property of RORγt was subsequently co-opted to support diverse immune cell functions in vertebrates.

### Limitations

Our *in vitro* studies used the W601 nucleosome positioning sequence, which is absent from native chromatin^61^. This was acceptable, because the stability of W601-based substrates is important for biochemistry and structural biology, and attempts to substitute endogenous sequences for structural studies have been marred by challenges^52^. Although our mononucleosomes include free DNA linkers, thereby representing the basic unit of chromatin, dinucleosomes and larger substrates may better reflect higher-order chromatin dynamics that TFs encounter^62^. Finally, without nucleosome maps for Th17 cells, we cannot place ROREs within endogenous chromatin, so our inferences rely on correlations between *in vitro* biochemistry and cellular function. Future efforts will address these limitations.

## ACKNOWLEDGEMENTS

We would like to thank Tony Hunter and Chris Glass for providing feedback and reviewing an early version of this manuscript. This work was supported by the NOMIS Foundation, The Morris Foundation, The National Institute of Health and National Science Foundation through grants F32 GM148049 and K99 GM159035 awarded to TSS, R01 DK133587 awarded to PRG, U01 AI136680, U54 AI170855, R01 GM151305 awarded to DL, R01 AI151123 awarded to YZ, and R01 AI196844 awarded to P.R.G., D.L., and Y.Z, R35 GM153502 awarded to P.C.W. D.L. also acknowledges support from the National Science Foundation grant MCB-2048095, NIH-NCI CCSG P30 CA014195, as well as support from the Margaret T. Morris Foundation and the Hearst Foundations. The authors acknowledge Dr. Laura Koeppling for assistance in collecting cryo-EM datasets at the Sanford Burnham Prebys Cancer Center cryo-EM center supported by NCI Cancer Center Support Grant P30 CA030199 and NIH Shared Instrumentation Grant S10 OD026926. Work at the Center for Theoretical Biological Physics was supported by the NSF (Grants PHY-2019745 and PHY-2607619).

## DATA AVAILABILITY

HDX-MS data are available as processed metadata and raw files. The proteomics data are available via ProteomeXchange with identifier PXD079146. Final electron density maps, half maps, local resolution data, filtered maps and pixel-size calibrated maps are available for RORγt-NCP and the NCP datasets at EMDB with identifiers EMD-77866 and EMD-77867, respectively. Atomic models of the RORγt-NCP and the NCP complexes have been deposited at the PDB and can be accessed using identifiers pdb_000036ug and pdb_000036uh, respectively. RNA-seq data are available as read count values and raw sequencing reads. CUT&RUN data are available as raw sequencing reads, alignments in bigwig format, and peak calls in narrowPeak format. RNA-seq and CUT&RUN data are available on the Gene Expression Omnibus with identifiers GSE334128 and GSE334124, respectively.

## MATERIALS AND METHODS

### Chemicals and reagents

Unless stated otherwise, all chemicals were purchased from Sigma Aldrich. Lyophilized human histones (H2A, H2B, H3.1, H4) were purchased from The Histone Source at Colorado State University. Tris-HCl and HEPES pH 7.5 purchased from Teknova. Guanidine hydrochloride was purchased from BioVision Inc. Dithiothreitol (DTT) purchased from Biopioneer Inc. Ammonium sulfate and TEMED purchased from Biorad. Bovine serum albumin (BSA) purchased from New England Biolabs. Potassium chloride and Chloroform:isoamyl alcohol were purchased from VWR. Sodium acetate was purchased from MP Biomedicals. Bis-acrylamide and Phenol:chloroform:isoamyl alcohol were purchased from Fischer Scientific. Magnesium sulfate purchased from Macron Fine Chemicals.

### RORγt expression and purification

We performed insertion PCR to introduce a SRC2 peptide on the C-terminus of the RORγt LBD in the pESUMO-RORγt construct (Addgene 170329) using the Q5 mutagenesis tools (NEB). This peptide artificially stabilizes the LBD in an active conformation in the absence of ligand ^54,63^. This was done to reduce the number of variables such as ligand binding and coregulator binding. This modification also improved the expression yield from ∼6 to ∼15 mg of protein per liter of expression media. HisSUMO-RORγt-SRC2 was expressed and purified using previously described methods ^34^ with some modifications listed below. BL21 (DE3) transformed with the pESUMO-RORγt-SRC2 plasmid was cultured in terrific broth supplemented with 50 μg/mL carbenicillin and 30 μM zinc chloride. The culture was grown with orbital shaking at 150 RPM at 37 °C to optical density of 0.5 at 610 nm, the temperature was reduced to 16°C, and then IPTG was added to 250 μM for overnight expression. The cells were harvested and washed with NiB1 (50 mM HEPEs pH 7.9, 500 mM NaCl, 25 mM imidazole, 10% glycerol) supplemented with 1X SigmaFast protease inhibitor cocktail (Sigma), and snap frozen in liquid nitrogen and stored at −80°C. The pellets were thawed and resuspended in lysis buffer (NiB1 supplemented with 1X SigmaFast protease inhibitor cocktail and Benzonase Nuclease from Sigma). The cells were lysed using sonication operating at 30% amplitude with 6 seconds on and 6 seconds off for a total of 5 minutes per cycle. We repeated the sonication 3-4 times until the sample was homogeneous. The lysates were clarified with high speed centrifugation at 30,000 relative centrifugal force (RCF). The supernatants were used for NiNTA affinity chromatography and ion exchange chromatography as previously described. After elution from the ion exchange column, the protein products were concentrated with amicon ultra centrifugation columns (Sigma) to 48 μM concentration, determined by absorbance at 280 nm using a NanoDrop nanospectrophotometer (Thermo Scientific). The protein was aliquoted, snap frozen, and stored at −80 °C. On the day of use, 50 μL HisSUMO-RORγt-SRC2 protein samples were thawed, mixed with 5 μL His-tagged SUMO protease (gifted by the Griffin Lab), and then size exclusion purified into reaction buffers appropriate for the analysis as described below. The S92A/L93A and K31 mutations were introduced to the pESUMO-RORgt-SRC2 plasmid using the Q5 mutagenesis tools (NEB). Mutant proteins were expressed and purified using the same procedure described above.

### Reconstitution and purification of human histone octamer

Lyophilized human histones H2A, H2B, H3.1, and H4 (purchased from The Histone Source at Colorado State University) were mixed at a molar ratio of 1:1:0.9:0.9, respectively, in denaturing buffer (20 mM Tris-HCl pH 7.6, 6 M guanidine hydrocholoride, and 5 mM dithiothreitol). The protein mixture was dialyzed against refolding buffer (10 mM Tris-HCl pH 7.6, 2 M NaCl, and 1 mM dithiothreitol). The dialysate was concentrated and purified via size-exclusion chromatography (Superdex 200 Increase 10/300; Cytiva) equilibrated in refolding buffer. The resulting octamer was concentrated by centrifugal filtration (Amicon Ultra, Millipore) and stored at 4°C.

### DNA preparation and purification

DNA for nucleosome assembly was generated by PCR amplification. Briefly, Pfu polymerase was expressed and purified, and the standard Pfu PCR conditions Pfu buffer (20 mM Tris-HCl pH 8.8, 10 mM ammonium sulfate, 10 mM KCl, 0.1 mg/ml BSA, 0.1% (v/v) Triton-X-100, 2 mM magnesium sulfate) was used to generate DNA at milligram scale. The resulting DNA fragments were extracted with phenol:chloroform:isoamyl alcohol mixtures using the MaXtract High Density gel column (Qiagen), followed by a chloroform:isoamyl extraction. DNA was extracted from the aqueous solution using sodium acetate ethanol extraction. The resulting DNA fragment was purified by anion-exchange chromatography (MonoQ 5/50 GL; Cytiva). The DNA was eluted from the MonoQ 5/50 GL column following a gradient of increasing salt concentration up to 1 M NaCl (0% NaCl for 20 ml, 20% NaCl for 10 ml, 20-60% NaCl for 15 ml, 60-80% NaCl for 22.5 ml, 80-100% NaCl for 1.5 ml, 100% NaCl for 3.5 ml). DNA fragments were extracted overnight at −20°C using sodium acetate ethanol extraction. The resulting DNA fragments were resuspended in 10mM Tris-HCl pH 8.0, assayed for size and purity by TapeStation (Agilent), and stored at 4°C for immediate use or −20°C for long-term storage.

### General nucleosome assembly and purification

Nucleosomes were reconstituted using a previously described method ^64^. All steps were performed on ice or at 4° C unless otherwise noted below. Briefly, DNA and octamer were mixed in high salt reconstitution buffer (10 mM HEPES or 10 mM Tris-HCl pH 7.5, 2 M KCl, and 1 mM dithiothreitol). DNA was held at a constant concentration (typically 3 μM or 6 μM) and the ideal molar ratio of histone octamer was determined empirically by testing molar ratios between 0.8 and 1.2 times the concentration of the DNA. The DNA octamer mixture was initially dialyzed against high salt reconstitution buffer for 30 minutes in Slide-A-Lyzer MINI 3,500 MWCO dialysis button devices (Thermo Scientific). The KCl concentration was gradually reduced from 2 M to 0.25 M over a 16 hour gradient by pumping low salt reconstitution buffer (10 mM HEPES or Tris-HCl pH 7.5, 0.25 M KCl, and 1mM dithiothreitol) using two peristaltic pumps. The samples were dialyzed against fresh low salt reconstitution buffer for three hours, followed by dialysis against nucleosome storage buffer (10mM HEPES or Tris-HCl pH 7.5, 1 mM DTT) for three hours, and then dialyzed against fresh nucleosome storage buffer overnight. After dialysis, the quality of the nucleosomes was assessed by native polyacrylamide gel electrophoresis using 5-6% PAGE gels (59:1 acrylamide:bis-acrylamide). Native polyacrylamide gels were prepared by combining the PAGE gel solution described previously, 10% ammonium persulfate, and TEMED in a gel cassette (1.0 mm Novex), followed by overnight polymerization. Once the ideal ratio of DNA and octamer was determined, nucleosome assembly reactions were scaled up and dialyzed using the method described above. After dialysis, nucleosomes were incubated at 37 °C for thirty minutes before being purified by native polyacrylamide gel (7%) electrophoresis using the Prep Cell 491 apparatus (Bio-Rad) in nucleosome storage buffer (10W, 2-7 hour run time). The eluted nucleosomes were concentrated with an Amicon Ultra centrifugal filter (Millipore) and were stored in nucleosome storage buffer on ice until use.

### Selected Engagement of Nucleosomes coupled with Sequencing (SeEN-seq)

#### SeEN-seq Assay Development

We adapted a protocol based on the originally described SeEN-seq method ^32^, overviewed in Fig. 1a. Briefly, we elected to use a nucleosome positioning sequence with 30 bp extensions to allow for RORγt to bind to free DNA. The 30 bp extensions also contained 15 bp priming sites that allow for amplification and functionalization of the library. This created a 177 bp tiling region across the Widom W601 nucleosome positioning sequence ^65^, as shown in Extended Data Fig. 1a. We tiled two known 11 bp ROREs through the tiling region, yielding a library size of 336 members. A cRORE based on a sequence from IL-17 enhancer element ^33^ and a vRORE based on a sequence from the androgen promotor ^10^ were used. We amplified and sequenced the library to examine the distribution of distribution of reads across all library members, Extended Data Fig. 1b. The FAM-labeled library was mixed with an unlabeled 207bp W601 sequence that did not contain priming sequences in a 1-to-1000 molar ratio. We used the DNA mixture for standard nucleosome core particle assembly and purification using established methods^64^. We performed selections by reacting the nucleosome samples with RORγt or buffer and separated reaction products from reactants using electrophoretic mobility shift assay, Extended Data Fig. 1c. Using fluorescence scanning, we cut out small sections of the gel containing the shifted band as well as the same region of the gel in the buffer treated negative control sample. We extracted DNA and performed quantitative PCR to ensure an enrichment of in the RORγt treated sample compared to the negative control, Extended Data Fig. 1d. We amplified and sequenced the library after one round of selection to identify the sequences that had been enriched based on changes relative abundance, Extended Data Fig. 1e. We normalized the relative read counts to the input library to calculate the fold change in enrichment, Extended Data Fig. 1f. We investigated performing a second round of selection by repeating the process and including additional selection replicates, Extended Data Fig. 1g. The qPCR trace showed a similar signal-to-noise compared to the first round of selection, Extended Data Fig. 1h. We also included an additional buffer only negative control which showed the background signal in qPCR originating from the gel extraction process. After sequencing the library after the second selection, we observed that there was an increased signal for some of the library members and a decrease in signal for the non-binding sequences, Extended Data Fig. 1i. After normalizing the library abundances, the enrichment ratios for productive binding sequences increased while the non-binding sequences decreased, Extended Data Fig. 1h. Overall, the trends for one round and two rounds of selection are very similar. We conclude that a second round of selection may benefit signal-to-noise and produce more confident results although a single round of selection is likely satisfactory for simple cases.

#### DNA synthesis and amplification

SeEN-seq libraries were designed and synthesized using the Twist Bioscience Oligopool system. SeEN-seq libraries were PCR amplified using the KAPA HiFi polymerase kit (Roche) according to manufacturer’s instructions. Briefly, test reactions were first conducted using quantitative PCR by including EvaGreen dye (Biotium) and following fluorescence using QuantStudio 6 (ThermoFisher Scientific) to identify suitable reaction conditions that did not over cycle the library. We then performed a larger reaction to generate ∼3 μg of library DNA. To fluorescently label the library DNA, FAM-labeled primers were used to amplify library DNA using low cycle (4-8 cycles) PCR. PCR products were cleaned up using AMPure XP (Beckman Coulter), assayed for size and quality by TapeStation (Agilent), quantified using Qubit assay (Thermo Fisher), and stored at 4°C for immediate use or −20°C for long-term storage.

#### Selection Assay

The FAM-labeled library was mixed with an unlabeled 207bp W601 sequence that did not contain priming sequences in a 1-to-1000 molar ratio. We used the DNA mixture for standard nucleosome assembly and purification as described above. We performed selections by reacting the 1 μM nucleosome samples with 500 nM RORγt or buffer only control at 25°C for 30 minutes. The reaction buffer contained 20 mM HEPES pH 7.9, 100 mM NaCl, 5% glycerol, 1 mM DTT. and separated reaction products from reactants using electrophoretic mobility shift assay. We found that 5% PAGE gels (59:1 acrylamide:bis-acrylamide) were able to separate RORγt bound nucleosomes from unbound nucleosomes. We performed fluorescence imaging with the Amersham Typhoon (Cytiva) to isolate signal from the SeEN-seq library DNA. We cut out the bands and performed crush soak gel extraction by incubating the bands overnight in 50 mM Tris pH 8.3, 300 mM NaCl, 0.05% Tween-20, 1 mM EDTA. The gel extract was syringe filtered at 0.22 um and then used for overnight sodium acetate ethanol extraction with GlycoBlue coprecipitant (Thermo Fisher). The precipitates were isolated with centrifugation, washed with freshly prepared 70% ethanol, dried, and then resuspended in storage buffer (5 mM Tris pH 8.5). The DNA was amplified using the methods described above.

#### Next-generation sequencing and data analysis

We sequenced the unlabeled SeEN-seq library DNA samples using the Amplicon-EZ service at Genewiz. Briefly, 500 ng of material was prepared for paired-end 250 bp sequencing reaction through a proprietary Illumina-based sequencing platform (Azenta Life Science). Reads were aligned to the designed library sequences using Bowtie2 (version 2.4.4) with options `--no-discordant --no-mixed --local --score-min L,0,2 --minins 205` –`maxins 207`. We found it was important to include a stringent read scoring parameters improved the alignment results. The scoring scheme allows for a single mismatch or 2 terminal base truncation, and does not tolerate insertions. The alignments rates ranged from 20-40% of total reads. Read counts per member of the library were extracted using `idxstats` (samtools version 1.13). Read count data was imported into Rstudio (version 2024.04.2+764) for normalization to total read counts (per sample) statistical analysis, comparison of selection rounds, and plotting.

### RORγt:Nucleosome Complex Assembly

The sequence for 207 bp W601 with cRORE at site 14 (superhelical location -6.5) was synthesized and cloned into the pUCIDT vector by Integrated DNA Technologies (IDT). The DNA was amplified using the SeEN-seq library primers as detailed in the general DNA amplification and purification protocol described above. We assembled and purified nucleosomes using the standard protocol described above. 50 μL aliquots of HisSUMO-RORγ-SRC2 (48 μM) was thawed and mixed with 5 μL HisSUMO protease (50 μL) and incubated on ice for 30 minutes. We optimized stoichiometries of nucleosome to RORγt using EMSA. Briefly, nucleosomes were mixed with excess RORγt-SRC2 at different molar ratios, reacted at 25°C for 30 minutes in 20 mM HEPES pH 7.9, 100 mM NaCl, 5% glycerol, 1 mM DTT. The reactions were assayed using 5% PAGE. We also performed analytical size exclusion using a 3.2/300 Superose 6 (Cytiva) on our AKTA pure FPLC (Cytiva) with running buffer containing 10 mM HEPEs pH 7.5, 50 mM NaCl, 1 mM DTT, and 5% glycerol. The fractions with native 5% PAGE and denaturing 4-20% PAGE. Stoichiometric ratios for complex assembly were optimized empirically by gel shift and size exclusion chromatography (SEC). A 3-to-1 molar ratio of RORγt to nucleosome effectively shifted the nucleosome band, such that the binary complex could be efficiently purified by SEC (Extended Data Fig. 2a). Denaturing PAGE stained with Coomassie confirmed that RORγt and histones comigrated (Extended Data Fig. 2b), and native PAGE stained for DNA confirmed that the complex was intact (Extended Data Fig. 2c). The cRORE was selected over the vRORE because of its higher SeEN-seq enrichment ratio, which implies higher RORγt binding affinity.

### Hydrogen-deuterium exchange mass spectrometry

#### Sample Preparation

HisSUMO-RORγ-SRC2 (500 μL, 48 μM) was mixed 1:10 with HisSUMO protease (50 μL) and incubated on ice for 30 minutes. Cleaved protein was separated using a size-exclusion Superdex 200 Increase 10/300 GL column (Cytiva, 28990944) pre-equilibrated with 2x reaction buffer (20 mM HEPES, pH 8, 200 mM NaCl, 2 mM DTT) on an AKTA pure (Cytiva, Marlborough, MA). Fractions were collected and run on denaturing SDS-PAGE gel (Bio-RAD, 4569036) and stained with Coomassie blue to identify correct fractions. Fractions were pooled and concentrated to 30 μM using a 30 kD MWCO spin column (Sigma, UFC5030). Protein concentrations were estimated using nanodrop A280 values and molar absorptivity from ProtParam^66^. Ligand 25-hydroxycholesterol was added to protein mixture to a final concentration of 40 μM. Cleaved RORγ-SRC2 was mixed 1-to-1 with nucleosome to yield 50 μL protein mixture, containing 15 μM RORγ-SRC2, 5 μM nucleosome, and 20 μM 25-hydroxycholesterol in a final buffer of 15 mM HEPES, pH 8.0, 100 mM NaCl, and 1 mM DTT.

#### Peptide identification

To generate a peptide set, peptides were acquired using MS/MS experiments performed on a QExactive (ThermoFisher Scientific, San Jose, CA) over a 70-min gradient. Product ion spectra were acquired in data-dependent mode, with the five most abundant ions selected for MS2. For peptide identification, the MS/MS *.raw data files were analyzed on Sequest (version 2.3, Matrix Science, London, UK). Mass tolerances were set to ± 0.6 Da for precursor ions and ± 10 ppm for fragment ions. Oxidation to methionine was set as a variable modification. Non-specific digestion was selected in the search parameters with 4 missed cleavages. Only peptides with a false discovery rate (FDR) <1% were used in the dataset, and all identified charge states were included for each peptide in the peptide set.

#### Continuous Labeling

Experiments were carried out on a fully automated system (CTC HTS PAL, LEAP Technologies, Carrboro, NC; housed inside a 4°C cabinet) as previously described ^67^ with the following modifications: RORγ-SRC2 and nucleosome were mixed as described in *Sample Preparation* then incubated at 25°C in a thermocycler for 30 minutes. The solutions were cooled to 4°C, then transferred to the automated system for analysis. The reactions (5 μL) were mixed with 20 μL deuterated buffer (15 mM HEPES, pD 8.3, 100 mM NaCl, and 1 mM DTT) then incubated at 4°C for 10, 30, 60, 300, 900, or 3600 s. A non-deuterated control was included as a baseline (t=0s). Following deuterium exchange, deuterated solutions were quenched with an acidic quench solution (5 M Urea, 50 mM TCEP, 1% TFA, pH 2.5) before on-line digestion and data acquisition.

#### HDX-MS acquisition

Samples were digested using an immobilized Fungal XIII/pepsin column (1-to-1 ratio, prepared in-house) at 50 μL/min flow rate (0.1% (v/v) TFA at 4°C). Digested peptides were trapped and desalted using a 2x10mm C8 trap column (Hypersil Gold, ThermoFisher Scientific). Trapped peptides were gradient eluted [4 to 40% (v/v) CH3CN, 0.3% (v/v) formic acid] on a 2.1x50mm C18 analytical column (Hypersil Gold, ThermoFisher Scientific) over 5 minutes. All acquisition steps were performed at 4°C. Eluted peptides were directly ionized via electrospray ionization, coupled to a QExactive mass spectrometer (ThermoFisher Scientific).

#### Data Rendering

The intensity-weighted mean m/z centroid value of each peptide envelope was calculated and converted into a percentage of deuterium incorporation. This is calculated by determining the observed averages of the undeuterated (t=0 s) and fully deuterated spectra using the conventional formula described elsewhere ^68^. The fully deuterated control, 100% deuterium incorporation, was calculated theoretically, and a 70% deuterium recovery was set to correct for back-exchange, accounting for the 80% final deuterium concentration in the sample (1:5 dilution in deuterated buffer). Statistical significance for the differential HDX data is determined by an unpaired t-test for each time point, a procedure that is integrated into the HDX Workbench software ^69^.

The HDX data from all overlapping peptides were consolidated to individual amino acid values using a residue averaging approach. For each residue, the deuterium incorporation values and peptide lengths from all overlapping peptides were assembled. A weighting function weights shorter peptides more than longer peptides. Weighted deuterium incorporation values were then averaged to produce a single value for each amino acid. The initial two residues of each peptide, as well as prolines, were omitted from the calculations. This approach is similar to one previously described ^70^.

Deuterium uptake for each peptide is calculated as the average of %D for all on-exchange time points, and the difference in average %D values between the unbound and bound samples is presented as a heatmap with a color code given at the bottom of each figure (warm colors for increased deuterium exchange [deprotection] and cool colors for decreased deuterium exchange [protection]). Peptides are colored by the software to display significant differences, determined either by a greater than 5% difference (less or more protection) in average deuterium uptake between the two states or by using the results of unpaired t-tests at each time point (P < 0.05 for any two time points or P < 0.01 for any single time point). Gray peptides represent nonsignificant changes in the differential experiment (i.e., within ± 5% differential deuterium incorporation). The exchange of the first two residues for any given peptide is not colored. Each peptide bar in the heatmap view displays the average Δ %D values, associated SD, and the charge state. In addition, overlapping peptides with a similar protection differential deuterium exchange trends covering the same region are used to rule out data ambiguity. Output from data rendering was transferred to ChimeraX (version 1.8) to create protein structure overlays.

### Cryogenic Electron Microscopy (cryo-EM)

#### Graphene grid preparation

We adapted a protocol for preparing monolayer graphene grids that has been previously described ^71^. Briefly, we purchased Trivial Transfer® poly methyl methylacrylate (PMMA)-coated graphene pads from ACS materials and UltrAufoil R1.2/1.3 300 mesh grids from Quantifoil. We washed the grids by sequentially soaking them in chloroform, acetone, and isopropanol in 20 minute periods on an orbital shaker. After drying the grids, we plasma cleaned the grids using a Solarus II Model 955 (Gatan) plasma cleaning instrument. The grids were immediately submerged into a water bath and PMMA-coated graphene was floated onto the grids using the manufacturer’s protocol. We recovered the PMMA-coated graphene grids, allowed them to dry for 20 minutes at room temperature, and then baked the grids at 300 °C for 60 minutes. After letting the grids cool to room temperature, we submerged them in acetone for 60 minutes. We transferred the grids to a clean acetone solution and repeated the acetone wash this time overnight. The next day, the acetone was exchanged for a third wash for 60 minutes, then a subsequent wash with isopropanol for 30 minutes. All washes of the grids were performed in clean crystallization dishes covered in aluminum foil, on an orbital shaker in a fume hood at room temperature with gentle agitation. The graphene grids were dried at room temperature and then baked at 200 °C for 20 minutes. Grids were then either immediately used or stored in grid boxes and vacuum sealed.

#### Sample preparation for cryo-EM

The RORγt:nucleosome complex assembly was performed as described above with some minor modifications. We reacted 2 μM of nucleosomes with 6 μM of RORγt and SUMO protease in standard reaction buffer. After complex assembly reaction was completed, glutaraldehyde (Electron micrscopy sciences) was added to a final concentration of 0.01%. The complex was crosslinked for 10 minutes on ice, and then quenched by the addition of Tris pH 8.0 to a final concentration of 50 mM. The crosslinked sample was then purified with analytical size exclusion using a 3.2/300 Superose 6 (Cytiva) on our AKTA pure FPLC (Cytiva) with running buffer containing 10 mM HEPEs pH 7.5, 50 mM NaCl, 1 mM DTT. The material eluted at ∼300 nM based on absorbance at 260 nm which was found to be ideal for achieving good particle distribution on graphene grids.

#### Vitrification

Graphene grids were plasma cleaned using a Solarus II Model 955 (Gatan) plasma cleaning instrument. Grids were immediately loaded into a vitrobot mark IV (thermos fisher) operating at 4 °C and 100% humidity. 2 μL of ∼300 nM RORγt:nucleosome complex in size exclusion buffer was applied to the graphene surface of the grid, incubated for 30 seconds, and then 2 μL of size exclusion buffer + 0.02% DDM was added to the specimen. The sample was immediately blotted and plunge frozen in liquid ethane. We repeated this process with ∼300 nM of crosslinked nucleosome sample. The grids were clipped, screened, and ultimately used for data collection.

#### Data Collection

All data collection statistics are summarized in Extended Data Table 2. For RORγt-Nucleosome, we collected data on an FEI Titan Krios (Thermo Fisher Scientific) equipped with a K3 direct electron detector (Gatan). The data collection was automated using serialEM ^72^. We collected 9668 movies with a total dose of 49.23 electrons/Å^2^ at a rate of 0.7 electrons/pixel/frame at a pixel size of 1.06 Å/pixel. We collected 72 frames per movie, with a random defocus ranging from −0.7 to −2.5 um during collection. For Nucleosome only, we collected data on a Titan Krios G4 (Thermo Fisher Scientific) equipped with a Falcon4i detector (Thermo Fisher Scientific). The data collection was automated using EPU (Thermo Fisher Scientific). We collected 4058 movies with a total dose of 50.0 electrons/Å^2^ at a rate of 0.83 electrons/pixel/frame at a pixel size of 0.954 Å/pixel. We collected 60 frames per movie, with a random defocus ranging from −0.7 to −2.5 μm during collection.

#### Data Processing of RORγt:nucleosome

The overview of our processing scheme is shown in Extended Data Fig. 3. Briefly, CryoSPARC version 4.6.2 was used for preprocessing, 2D classification, and *ab initio* 3D reconstruction Extended Data Fig. 3A. We used Relion version 5.0b to perform 3D classifications, Extended Data Fig. 3e. Raw movies and the gain reference were loaded into CryoSPARC version 4.6.2 ^73^. The data was preprocessed using patch motion correction and patch CTF estimation, an example is shown in Extended Data Fig. 3b. We performed two rounds of particle picking starting with initial template matching and convolutional neural network approach with Topaz ^74^ in parallel. We pooled the particle picks, removed redundant picks from the two approaches, extracted and 2D classified. The initial 2D classes showed a lot of features consistent with free DNA, Extended Data Fig. 3c. The particles that resolved into clear nucleosome particles were used to retrain the convolutional neural network and a second round of picking with templates and Topaz. We again pooled the particle picks, removed redundant picks, extracted, 2D classified, and removed bad particle picks. Representative 2D classes after optimizing the particle picking procedure are shown in Extended Data Fig. 3d. In the end, there were ∼1.41 million particle picks that were used for *ab initio* reconstruction. We performed nonuniform refinement and imposed C2 symmetry to align particles to the pseudo C2 symmetry axis that runs through the dyad of the nucleosome. Subsequently, the particle stack was imported into Relion, symmetry expanded, and used for focused 3D classification using the masking strategy show in Extended Data Fig. 4a. Important parameters for this process were to disable alignment, use blush regularization^75^, and to adjust the regularization parameter T to 50. After 3 rounds of 3D classification without alignment, any potentially duplicated particles were removed. The final particle stack was imported into CryoSPARC for final reconstruction, gold standard global Fourier shell correlation (FSC) analysis (Extended Data Fig. 4a), 3D FSC analysis (Extended Data Fig. 4b), and sampling compensation factor calculation ^76,77^, Extended Data Fig. 4c. The final map is shown in Extended Data Fig. 4d-e. The pixel size of final maps was manually adjusted to maximize correlation between the experimental map and a high resolution crystal structure model of the nucleosome (PDB ID 1KX5).

### Integrative Modeling of RORγt:nucleosome

We used Alphafold3^46^ to predict structures of RORγt bound to the nucleosome at SHL −6.5. Alphafold3 was able to predict the RORγt DBD:RORE interaction at the correct sequence in every prediction. Alphafold3 generally predicted that the LBD would interact with the nucleosome, but the position and orientation was strikingly different for each prediction and the predicted alignment error for the LBD was > 30 Å. We examined the different maps for agreement to the experimentally determined structure and HDX-MS observations. We then used ISOLDE^78^ to relax the predicted structure into the experimental structure as implemented in ChimeraX version 1.8^79^. Afterwards, we performed high resolution modeling using iterative refinement with Phenix v1.21^80^ and manual model fitting with winCoot v1.3.1^81^. Final model statistics were calculated using MolProbity web server^82^.

### Electrophoretic Mobility Shift Assay

FAM-labeled 207bp W601 with cRORE at site 14 (SHL −6.5) was amplified using FAM-labeled SeEN-seq library primers. The FAM-labeled DNA was mixed with an unlabeled 207bp W601 sequence that did not contain priming sequences in a 1-to-1000 molar ratio. We used the DNA mixture for standard nucleosome core particle assembly and purification as described above. We reacted 1 μM nucleosome samples with a range of RORγt-SRC2 concentrations and included a buffer only control at 25°C for 30 minutes. The reaction buffer contained 20 mM HEPES pH 7.9, 100 mM NaCl, 5% glycerol, 1 mM DTT. and separated reaction products from reactants using electrophoretic mobility shift assay. We found that 5% PAGE gels (59:1 acrylamide:bis-acrylamide) were able to separate RORγt bound nucleosomes from unbound nucleosomes. We performed fluorescence imaging with the Amersham Typhoon (Cytiva) to isolate signal from the SeEN-seq library DNA.

### Molecular Dynamics Simulations

Explicit-solvent simulations were performed using the cryo-EM acquired structure of the DBD of RORγt (residues I8-V97) bound to cRORE (Chain J: residues C163-G180, Chain I: residues C28-G45). Simulation input files were generated with tleap (Ambertools)^83^, using the Amber-ff19SB^84^ force field for proteins and OL24^85^ force field for DNA. Two Zn^2+^ ions, parameterized with the Zinc AMBER Force Field (ZAFF)^86^, were added to the Cys4 zinc finger motifs. This was accomplished by aligning the current structure with a Zn-containing structure of RORγ (PDB ID: 7XVN)^30^. The system was solvated with OPC^87^ water molecules in a box with a 10 Å buffer. 26 Na^+^ ions were introduced to neutralize system system, with parameters described by Li and Merz^88^. During molecular dynamics simulations, position restraints were initially applied to the heavy atoms of the solute (protein/DNA). The L-BFGS^89^ algorithm was used for minimization with three successive force tolerances (1000,100 and 10 kJ/mol/nm). The system was equilibrated for 5 ns with the NVT ensemble (293 K), followed by 5 ns with NPT (293 K, 1atm) using the Langevin middle integrator^90^, where NPT simulations used the Monte Carlo Barostat^91^. Another round of L-BFGS minimization was applied with force of 10 kJ/mol/nm. Three replicas of production runs were performed for two μs each. For the mutant, S92 and L93 were each replaced with Ala, and the system was solvated in a truncated octahedron box with 13 Å of buffer. Minimization and simulations followed the same protocols as used for the WT. All simulations were performed using OpenMM^92^ software (version 8.2).

Analysis was performed using MDAnalysis^93^. Secondary structure calculated used the DSSP algorithm^94^. The angular deviation of CTE was calculated by first aligning each simulated frame to the cryoEM structure, based on the C-alpha atoms of residues I8-M85. Next, we defined the vector between the C-alpha atoms of S86 and V97 and calculated its angles with respect to that of the cryoEM structure.

### Retroviral Transduction of RORγt in *RORC* knockout CD4 T cells

#### CD4 T cell isolation and culture

Naive CD4+ T cells were isolated from the spleen and lymph nodes of 8-to 10-week-old C57/BL6 wild-type mice by negative selection using MojoSort™ Mouse CD4 T Cell Isolation Kit (BioLegend, Cat. No. 480033) with additional biotin beads anti-CD44 (clone: 1-M7), and anti-CD25(clone: PC61.5). These cells were seeded in 48-well plates pre-coated with rabbit anti-hamster antibodies and activated and cultured with 1μg/ml anti-CD3 (145-2C11; bioxcel), 1 μg/ml anti-CD28 (37.51; bioxcel), 2 ng/mL hTGF-β and 20 ng/mL mIL-6. Cells were cultured in complete RPMI-1640 medium containing 10% fetal bovine serum (FBS), 55 mM 2-mercaptoethanol, 1% Pen/Strep, 1X GlutaMax, 1 mM HEPES, 1 mM sodium pyruvate, and 1X non-essential amino acids at a density of 8 × 10⁵ cells/ml.

#### Retroviral Production

293T cells were cultured in 6 wells with DMEM supplemented with 10% FBS and 1% Pen/Strep at 70% confluency and transfected with 1.2 μg of MigR1-RORγt and 0.8 μg of the packaging vector pCL-Eco (Addgene, #12371) using Lipofectamine 3000 (Thermo Fisher, #L3000008), following the manufacturer’s protocol. The cell culture medium was refreshed 16 hours post-transfection. Retroviral-containing supernatants were collected at 48 and 72 hours post-transfection.

#### Spin-fection

For retroviral transduction, T cells were spin-infected with viral supernatants 16 hours post-activation (1,258 × g for 90 minutes at 32°C) in the presence of 5 μg/mL Polybrene (Millipore #TR-1003-G). The viral supernatant was replaced with the original culture medium after 3 hours of culture. A second retroviral transduction was performed 32 hours post-activation to achieve a high transduction rate. Transduced cells were cultured with TGFβ and IL-6 and harvested for analysis after 3 days of culture.

#### Cytokine Stimulation

Cells were treated with ionomycin (0.5μg/ml) and PMA (0.05μg/ml) at day 5 of activation for 1.5 hours. Subsequently, GolgiPlug (1μg/ml) was added, and the cells were incubated for an additional 3.5 hours before harvest.

#### Surface and Intracellular Cytokine Staining and FACS analysis

Antibody solutions were prepared in MAC buffer using the following concentrations: CD4-PreCP5.5 at 1:400, CD8-AmCyan (BV510) at 1:400, and Ghost Red-APC Cy7 at 1:1000. Cells obtained with/ without cytokine stimulation were incubated with antibody for 40 minutes at 4°C. For intra-cellular staining, the cells were fixed in Fixation/Permeabilization solution (BD Cytofix/Cytoperm) for 15 minutes and intracellular cytokine staining IL-17A-PE-Cy7 at 1:400 and RORγt-BV421 at 1:400 in permeabilization buffer. Flow cytometric analysis and sorting were performed on BD LSR II or BD FACS Aria Fusion supported by the Flow Cytometry Core Facility of the Salk Institute (RRID:SCR_014839) analyzed using FlowJo software.

### RNA Sequencing Analysis (RNA-seq)

GFP+ CD4+ live cells were sorted, and RNA was isolated following the protocol of the RNeasy Micro Kit (QIAGEN, Cat. No. 74004). RNA was submitted to Novogene Corporation for library preparation using poly A enrichment and sequencing in a 150 reaction paired end format on a NovaSeq X Plus Series instrument (Illumina). The paired RNA-seq reads were aligned to the GRCm38 reference genome using v2.5.3a of the Spliced Transcripts Alignment to a Reference (STAR) aligner ^95^. The RemoveNoncanonical option of the outFilterIntronMotifs parameter was selected to filter out alignments that contained non-canonical junctions. Hypergeometric Optimization of Motif EnRichment (HOMER) v4.11.1 ^96^ was then used to quantify each sample’s RNA expression. Differential expression analysis was conducted with DESeq2 v1.36.0 ^97^. Differentially expressed genes were called at Log2 fold change > 1 (or < −1) and −Log10 adjusted P value > 10.

### Cleavage Under Targets and Release Using Nuclease (CUT&RUN)

#### Chemicals, Reagents, and Buffers

The plasmid encoding pAG/MNase was a gift from Steven Henikoff (Addgene plasmid # 123461). The pAG/MNase protein construct was expressed and purified from the JM101 strain of bacteria as previously described ^98^ and stored at −80°C until needed. Mouse anti-FLAG M2 antibody in 1 mg/mL solution (50% glycerol, 10 mM sodium phosphate, and 150 mM NaCl, pH 7.4) and 5% digitonin solutions were purchased from Sigma-Aldrich (St. Louis, Mo). The base CUT&RUN buffer was comprised of 20 mM HEPEs, 150 mM NaCl, 0.015% digitonin, 0.5 mM spermidine, 1X EDTA-free complete protease inhibitors (ROCHE).

#### Chromatin Extraction

T cell cultures were harvested and washed twice in ice cold 1XPBS, and 500,000 cells were seeded in V-bottom 96 well plates (Nunc MicroWell plates Thermo Scientific #249944). The cells were pelleted at 2000 RCF and the cells were decanted. The cells were resuspended in 200 μL of Antibody wash buffer solution (Base buffer + 2 mM EDTA), pelleted at 2000 RCF for 5 minutes, and the cells were decanted. This wash was performed 1 additional time (2 washes total). The cells were resuspended in 100 μL of antibody solution (FLAG antibody diluted 1:100 in base buffer + 2 mM EDTA), incubated for 1 hour on ice, pelleted at 2000 RCF for 5 minutes, and the cells were decanted. The cells were resuspended in 200 μL of base buffer solution, pelleted at 2000 RCF for 5 minutes, and the cells were decanted. This wash was performed 2 additional times (3 washes total). The cells were resuspended in 50 μL of pAG/MNase solution (pAG/MNase was diluted to 1X in base buffer), incubated for 1 hour at 4 °C, pelleted at 2000 RCF for 5 minutes, and the cells were decanted. The cells were resuspended in 200 μL of base buffer solution, pelleted at 2000 RCF for 5 minutes, and the cells were decanted. This wash was performed 2 additional times (3 washes total). The cells were resuspended in 100 μL of calcium solution (base buffer + 2 mM CaCl_2_) and incubated for 30 minutes on ice. 100 μL of stop solution (base buffer + 20 mM EDTA and 4 mM EGTA) was added and the solutions were incubated at 37 °C for 15 minutes. The cells were pelleted by centrifugation at 2000 RCF for 5 minutes and the supernatant was harvested.

#### DNA Extraction, NGS Library Preparation, and Sequencing

100 μL of DNA extraction buffer (30 mM Tris pH 8.0, 3 mM EDTA, 0.6 mg/mL RNase, 0.15 mg/mL proteinase K, 0.3% SDS) was added to chromatin samples, and the solutions were incubated at 55°C for 1 hour. 30 μL of 3 M Sodium Acetate pH 5.5 and 2 μL of GlycoBlue Co-precipitant (Thermo Scientific) was added to each reaction and mixed. Each reaction was ethanol precipitated by addition of 1 mL of −80 °C 200 proof ethanol and incubated at −20 °C overnight. DNA was pelleted by centrifugation at 16,000 RCF for 5 minutes, and pellets were decanted. The pellets were washed with 70% ethanol, pelleted by centrifugation at 16,000 RCF for 5 minutes, and pellets were decanted. Each pellet was air dried under vacuum to remove residual ethanol, resuspended with 50 μL of 5 mM Tris buffer, and stored at −20 °C until needed. The DNA samples were used for NGS library preparation using the NEBNext® Ultra™ II DNA Library Prep Kit for Illumina (New England Biolabs #E7645S). We followed the manufacturer’s protocols but included the following adaptations as recommended by Nan Liu, Zhejiang University. The heat denaturation step after end repair reaction was reduced from 65°C for 30 minutes to 50 °C for 1 hour. We performed adaptor ligation using 0.6 μM adaptors and USER digestion as recommended by the manufacturer. We performed a size selection step using SPRI beads (Beckman Coulter) to select DNA fragments between 150 and 1000 bases in length (primarily to remove larger genomic DNA fragments). Libraries were amplified by preparing PCRs as instructed by the manufacturer and adding EvaGreen (Biotium) to 1X. We performed PCR and followed dsDNA accumulation each cycle in QuantStudio 6 instrument (Thermo Scientific). We used the thermocycling program recommended by the manufacturer but reduced the extension time from 75 seconds to 10 seconds. We stopped the thermocycling program after an extension stage when dye intensity was increasing in log-phase which was usually around 12-15 cycles. We performed one more extension for 5 minutes. We performed a size selection of PCR products using SPRI beads (Beckman Coulter) to select DNA fragments between 150 and 1000 bases in length. Quality control analysis of NGS libraries was conducted using TapeStation 4150 (Agilent) and yield was determined using Qubit 4 instrument (Invitrogen). NGS libraries were pooled and submitted to Novogene Corporation to sequenced on a NovaSeq X Plus Series (Illumina) in a paired end 150 reaction format.

#### CUT&RUN Data Preprocessing and Differential Binding Analysis

Adaptor sequences were removed from reads using trim_galore version 0.6.10 and cutadapt version 5.0. Trimmed read qualtity was assessed using FastQC version v0.12.1. Trimmed reads were aligned to the mouse reference genome mm10 using bowtie2 version 2.5.4 with the following options `-p 48 --very-sensitive --local -I 30 -X 1000 --dovetail --no-mixed --no-discordant`. Poor quality reads were removed with Samtools view version 1.22.1 with the following options `-F780 -f2 -q30`. Resulting bam files were sorted and indexed with samtools. We merged bam files from biological replicates and converted to bigwig format with bamCoverage version 3.5.6 with the following settings `--binSize 10 --normalizeUsing RPGC --ignoreForNormalization chrX --extendReads 150`. We used igv-notebook version 0.6.2 implemented in a Jupyter notebook environment (jupyterlab version 4.4.5) to visualize alignments at selected genomic loci. For differential binding analysis, we called peaks for individual replicate bam files using MACS3 version 3.0.4 with the following settings `--nomodel --cutoff-analysis -f BAMPE -g mm -B -q 0.1`. We used DiffBind version 3.20.0 to perform quantitative analysis of our CUT&RUN datasets. We used a Trimmed Mean of M-values to normalize libraries for quantification.

### Chromatin Immunoprecipitation coupled with Sequencing (ChIP-seq) Data Analysis

Compared with CUT&RUN, ChIP-seq peaks are broader and contain signal from neighboring genomic contexts (Extended Data Fig. 8b); these datasets also contained more peaks, more reads and a higher fraction of reads within peaks (Extended Data Table 3). Reads for WT RORγt, S92A/L93A RORγt, and their empty vector (EV) negative control were downloaded from the sequencing read archive (Ascension code SRP104092) using SRA tool kit version 3.1.1. Adapter sequences were removed using TrimGalore version 0.6.10 and read quality was assessed before and after trimming using FastQC version 0.12.1. Trimmed reads were aligned to the mm10 mouse genome using bowtie version 1.3.1 with the following options `--chunkmbs 512 -p 48 -k 1 -m1 -v 2 --best –strata`. The resulting SAM files were filtered for junk alignments and converted to BAM files using samtools view (version 1.21) with the following options `-F 1804 -q 30`. Replicate BAM files were merged with samtools, converted to bedGraph format using bedtools version 2.31.1 and then to BigWig format for bedGraphToBigWig version 2.10. BigWig files were visualized using the integrative genome viewer for jupyter notebooks igv-notebook version 0.6.2.

### BPNet Analysis

#### Data Preprocessing

We converted merged and sorted BAM files from our CUT&RUN experiments and from previously published ChIP-seq experiments (described above) into sorted stranded tag-align bedgraph format with bedtools v2.31.1 genomecov with options `-5 -bg -strand <+/-> | sort -k1,1 -k2,2n`. We obtained a previously described ^99^ genome blacklist from the Kundaje lab and remove reads in from the blacklisted genomic regions using bedtools subtract. Last, tag align bedgraph files were converted to tagalign bigwig format with bedGraphToBigWig version 2.10. Peak files were generated from unmerged BAM replicates using MACS3 callpeak with options `--nomodel --cutoff-analysis -f BAMPE -g mm -B -q 0.05`. The peak summits found in blacklisted regions of the genome were then removed using bedtools subtract.

#### Model Training and Analysis

We used the BPNet model architecture and loss function that have been previously described ^49^. BPnet models were trained using an implementation in pytorch v2.71 called bpnet-lite (2020 © Jacob Schreiber). Our models were trained using default hyperparameters (8 convolutional layers, 64 filter channels, learning rate of 0.001, batch size of 64, 50 training epochs). Chromosomes 1 and 6 (containing key RORγt binding loci for Il17a, Il17f, Il1r1b, and Il23r) were omitted from the training dataset and were used for validation data. The training data consisted of all remaining chromosomes in the mouse genome. The input window perceptive field was set to 2114 bp. Model fitting, sequence attribution, negative control sequence identification, attribution analysis, and marginalization were done using bpnet-lite command-line. Sequences and tag-aligned reads were automatically extracted from peaks and used for model training using *fit*. GC-matched negative control sequences were also automatically identified using *negatives*. Predicted counts and profiles were generated and used for attribution analysis using *predict* and *attribution,* respectively. Motif discovery was done using modisco-lite v2.4.0. For marginalization experiments, we selected motifs from Jaspar database corresponding to RORγt (MA1151.1), BATF (MA0835.2), and IRF4 (MA1419.1). We then performed in silico mutagenesis to introduce a motif of interest at all possible positions and calculated the marginal increase in predicted score for models trained on either WT or S92A/L93A ChIP-seq datasets. Attribution analysis used DeepLIFT/SHAP, which assigns base-resolution importance scores explaining the sequence features that drive a model’s prediction; comparing WT and S92A/L93A models distinguishes sequences whose contribution is unchanged from those sensitive to the CTE mutation. Figures from our analyses were generated using the tangermeme suite of tools^51^.

---

**Extended Data Fig. 1:**
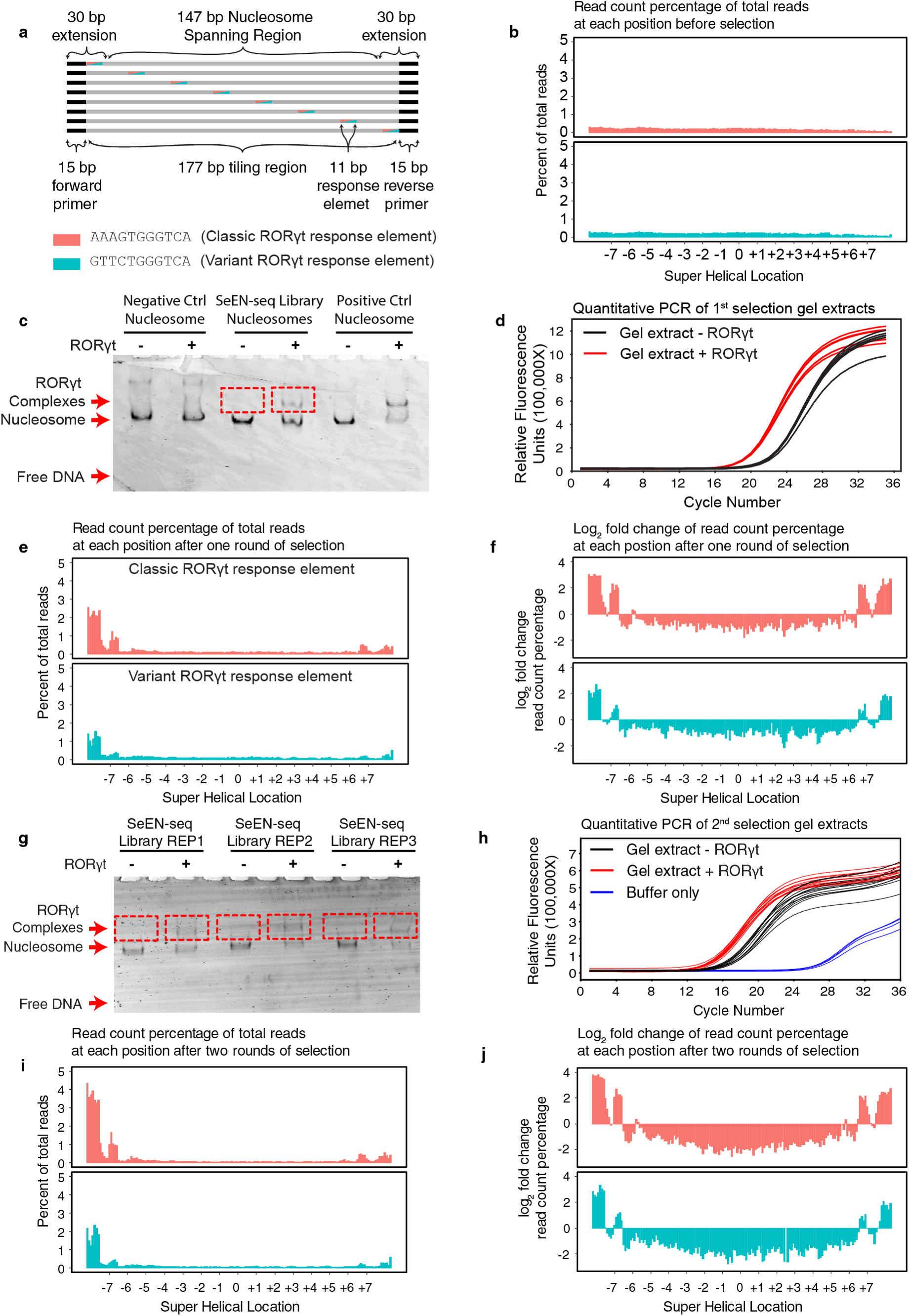
Development of SeEN-seq assay. (**a**) Schematic illustration of the library design. (**b**) The normalized read counts of the library members prior to selection. The read counts for each member are normalized by dividing by the total number of aligned reads and plotted as a function of SHL. (**c**) A representative gel image from the selection assay is shown. We included both positive and negative control nucleosomes for initial experiments to ensure that the selection assay promoted specific binding to ROREs, but did not promote nonspecific binding to nucleosomes that did not contain ROREs. The addition of the positive control nucleosome also helped identify where to cut the gel to extract the DNA (indicated by red boxes). (**d**) Quantitative PCR was performed on DNA extracted from the SeEN-seq library samples treated with RORγt and the buffer treated nucleosome. The DNA from the RORγt-treated samples was amplified and sent for sequencing. (**e**) The normalized read counts of the library members after one round of selection. (**f**) The log2 enrichment ratios of normalized read counts after one round of selection. (**g**) A representative gel image for the second round of selection. We performed the selection in triplicate for the second round and cut the bands as indicated by the red boxes. (**h**) Quantitative PCR of the extracted DNA. The DNA from the RORγt-treated was amplified and sent for sequencing. (**i-j**) As e-f, but after the 2^nd^ round of selection.

**Extended Data Fig. 2:**
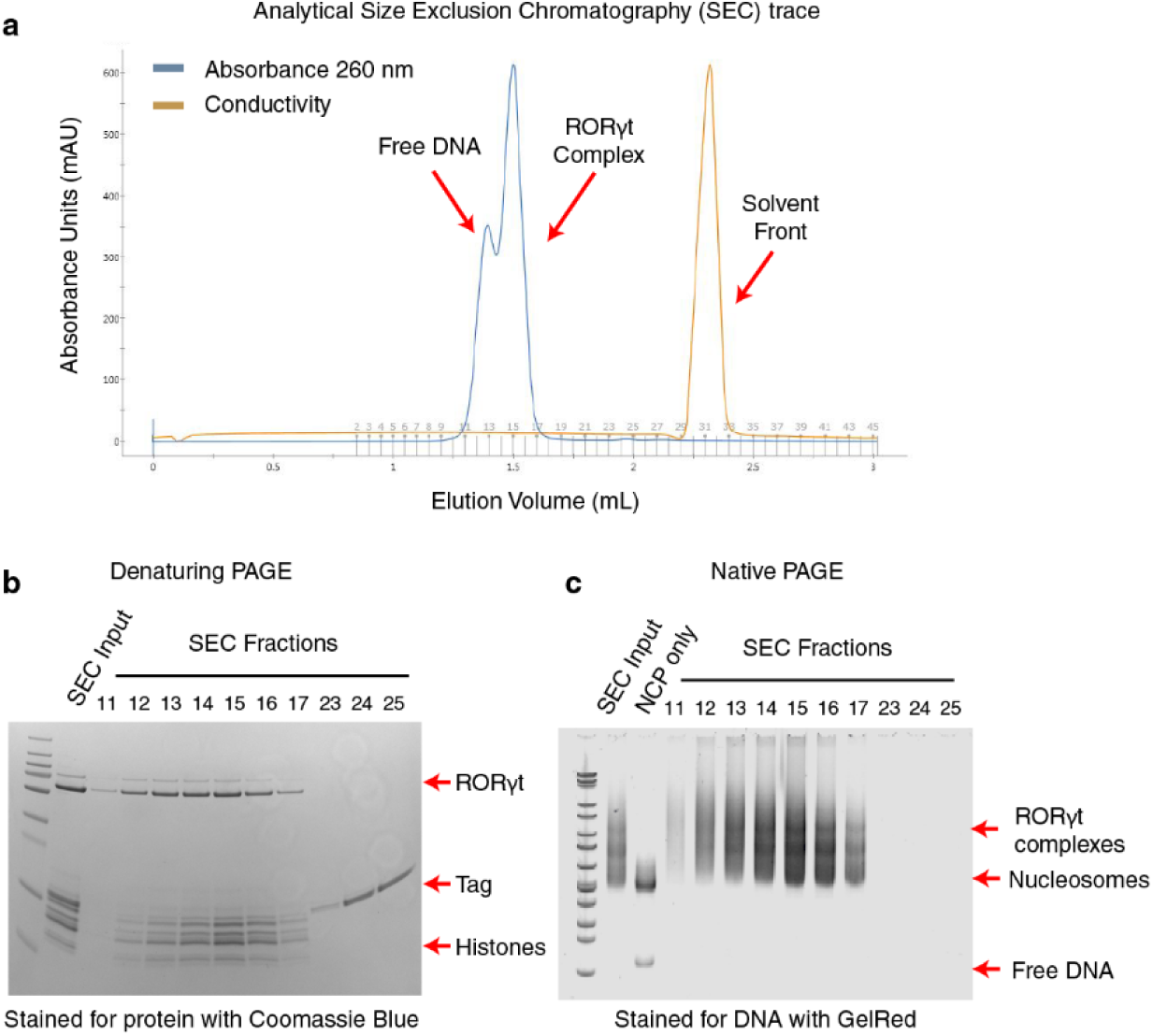
RORγt:nucleosome complex reconstitution. (**a**) RORγt and nucleosomes with cRORE installed at SHL −6.5 were used for complex assembly. A representative size exclusion chromatogram (SEC) of the complex purification. Selected fractions were assayed using (**b**) denaturing PAGE and (**c**) native PAGE.

**Extended Data Fig. 3:**
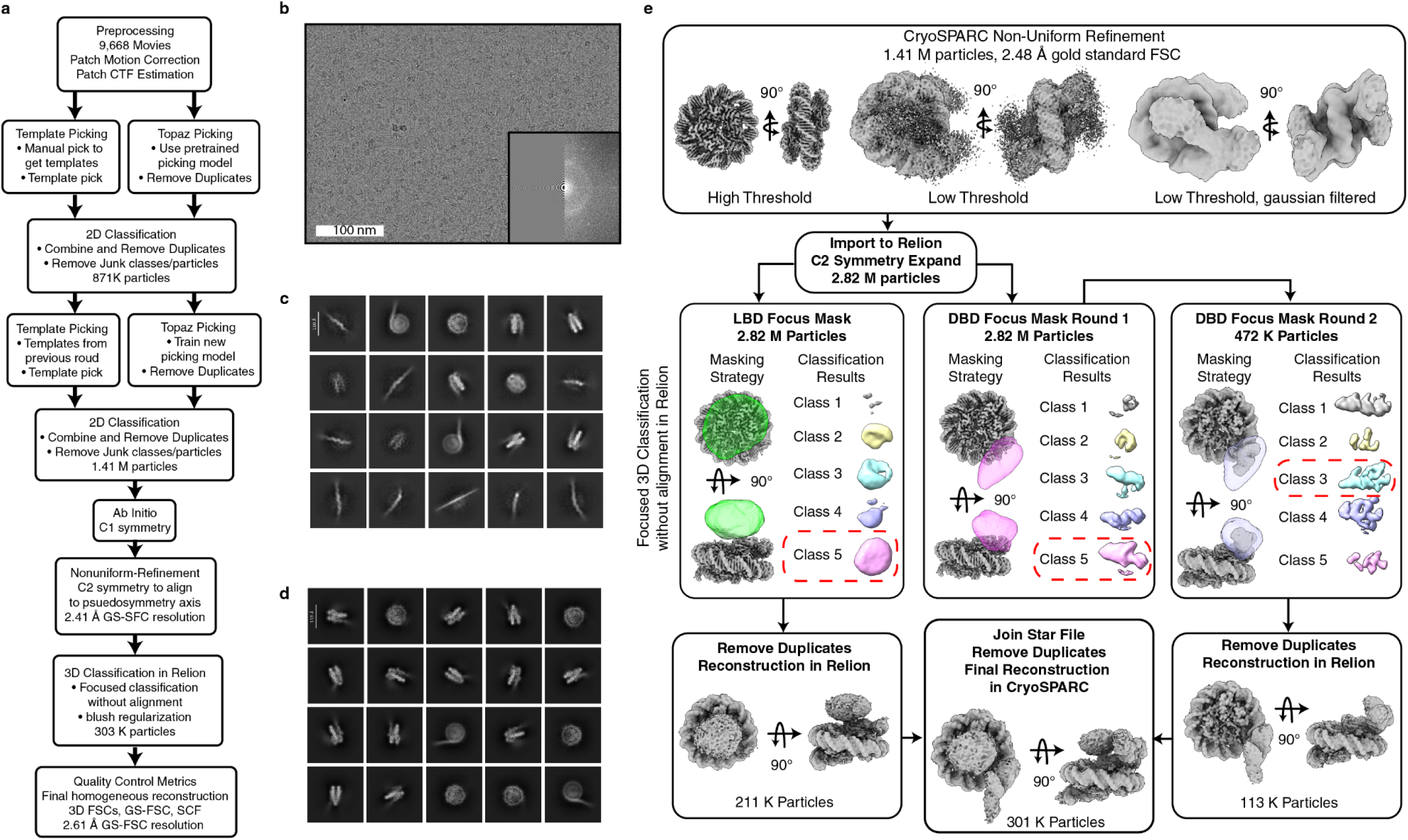
Cryo-EM data processing scheme and representative results. (**a**) The overall cryo-EM processing scheme for the dataset of RORγt engaged to SHL −6.5 on the nucleosome. All tasks were performed in CryoSPARC Version 4.6.2, unless otherwise noted. (**b**) Representative micrograph and 2D contrast transfer function (CTF) fit for the data shown alongside a power spectrum from the micrograph. (**c**) Results from an initial round of 2D classification. (**d**) Results from 2D classification after optimizing particle selection through Topaz. The scale bar in the 2D classification results is 110 Å. (**e**) The 3D classification strategy shown is implemented in Relion version 5.0b.

**Extended Data Fig. 4:**
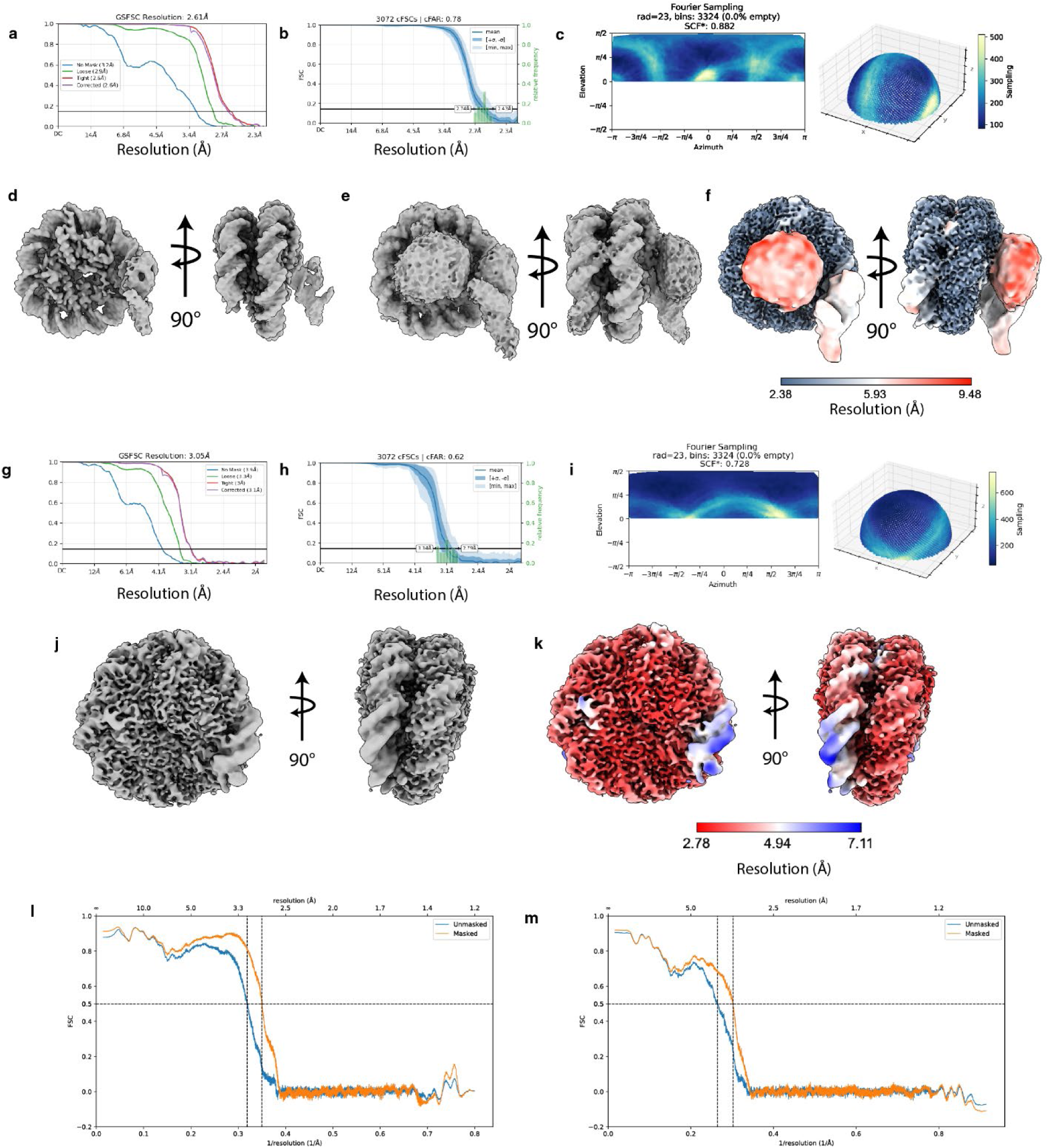
Validation of cryo-EM maps and models. Validation data is shown for RORγt-NCP complex in panels **a-f**. Validation data is shown for NCP only in panels **g-k**. (**a** and **g**) Gold standard Fourier Shell Correlation (FSC) analysis for the final map. (**b** and **h**) 3D FSC plots indicating the directional resolution of the reconstruction^100^. (**c** and **i**) The Fourier sampling distributions and sampling compensation factor calculations^76,77^ are shown. The final maps for the RORγt-NCP structure are shown in (**d**) at high threshold and in (**e**) at low threshold. The final map for the NCP only structure is shown in panel in (**j**). The final maps colored by local resolution are shown in panels (**f**) and (**k**). Map-to-model FSCs for the RORγt-NCP and NCP only structures are shown in (**l**) and (**m**), respectively.

**Extended Data Fig. 5:**
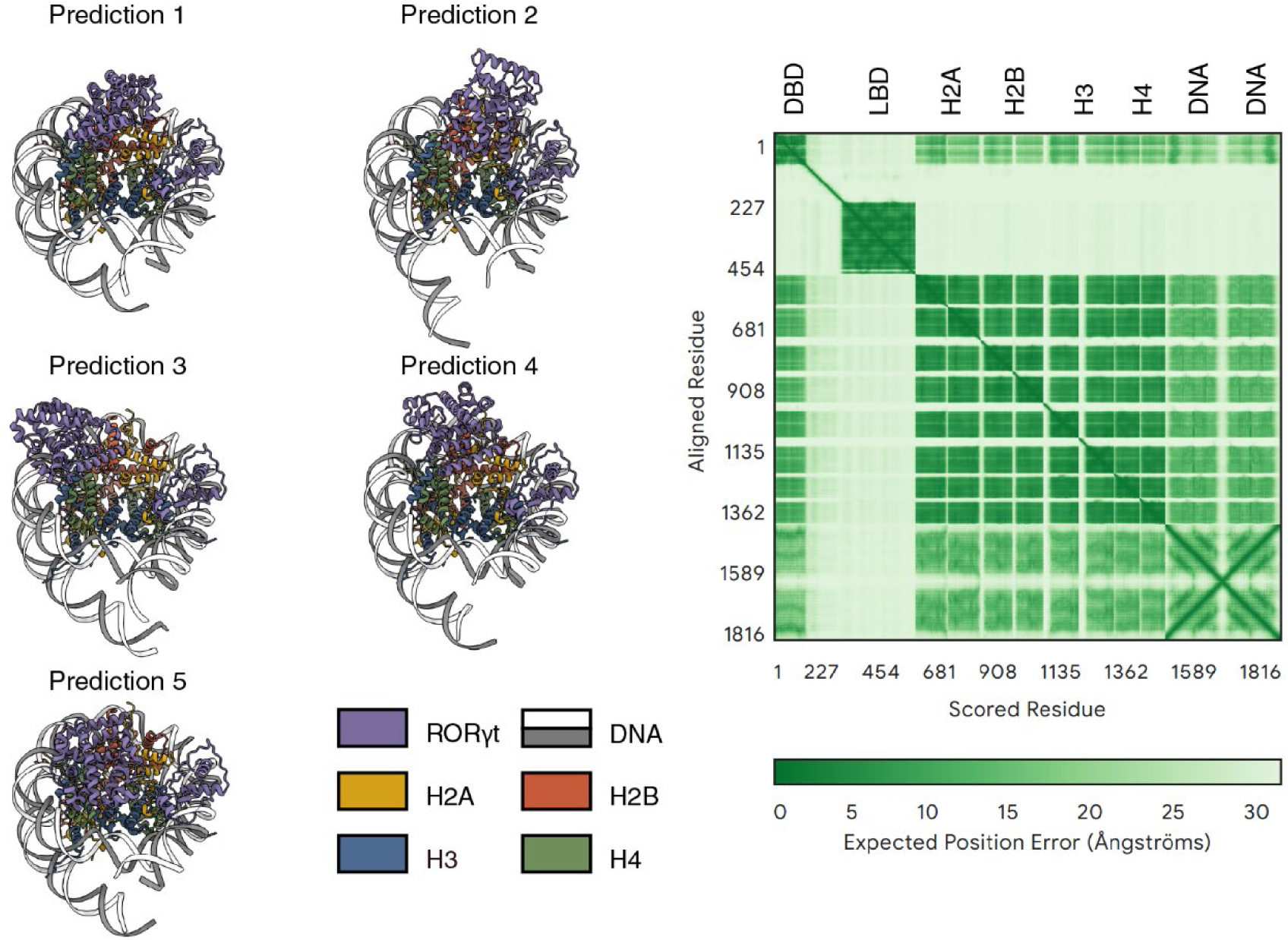
Representative Alphafold3 Structure Predictions. Alphafold predictions for the RORγt nucleosome complex are shown with histone tails, RORγt hinge region, and the DNA extensions hidden for clarity. A representative position alignment error plot is shown in the bottom right panel.

**Extended Data Figure 6:**
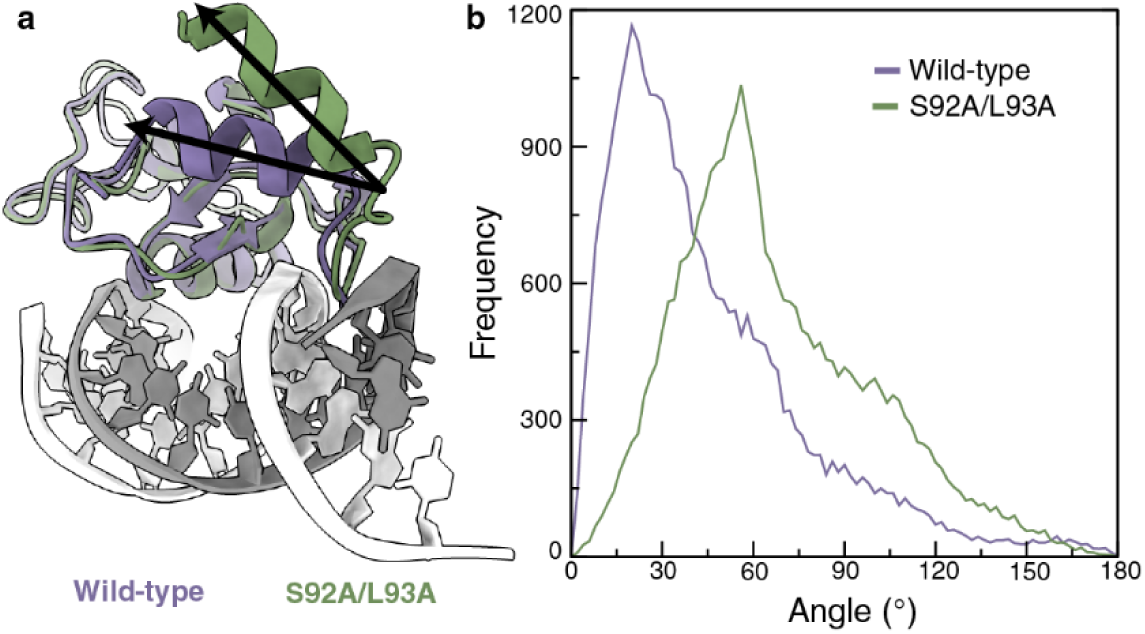
S92A/L93A mutations reposition the CTE relative to the DBD core. (**a**) Structural snapshot from an MD trajectory (green) overlaid with the experimental cryo-EM structure (purple), showing an angular deviation of the CTE. The helical parameters of the S92A/L93A CTE residues have also deviated from an idealized α-helix, which is apparent as a narrower helix in the cartoon representation. (**b**) Distribution of CTE angles for the WT and mutant proteins, calculated from MD simulations.

**Extended Data Fig. 7:**
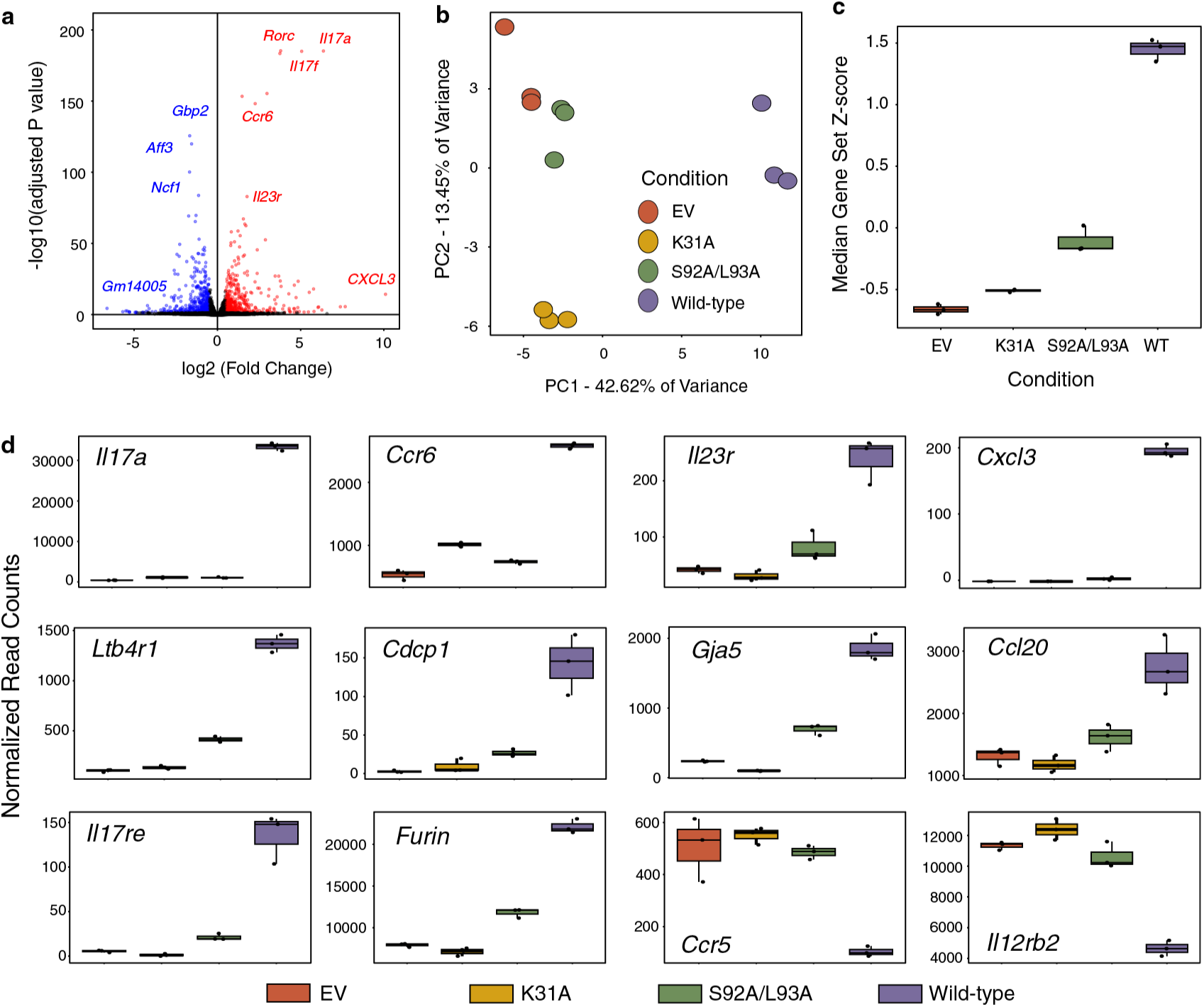
Genomic occupancy and transcriptomic analysis. (**a**) Differential RNA sequencing (RNA-seq) comparing the WT RORγt overexpression and the empty vector negative control. (**b**) Principal component analysis of RNA-seq datasets. (**c**) Median expression Z-score of the selected 27 gene dataset is plotted as a box plot. (**d**) Normalized read counts for selected genes are shown as box plots. Boxes are colored according to color key below.

**Extended Data Fig. 8:**
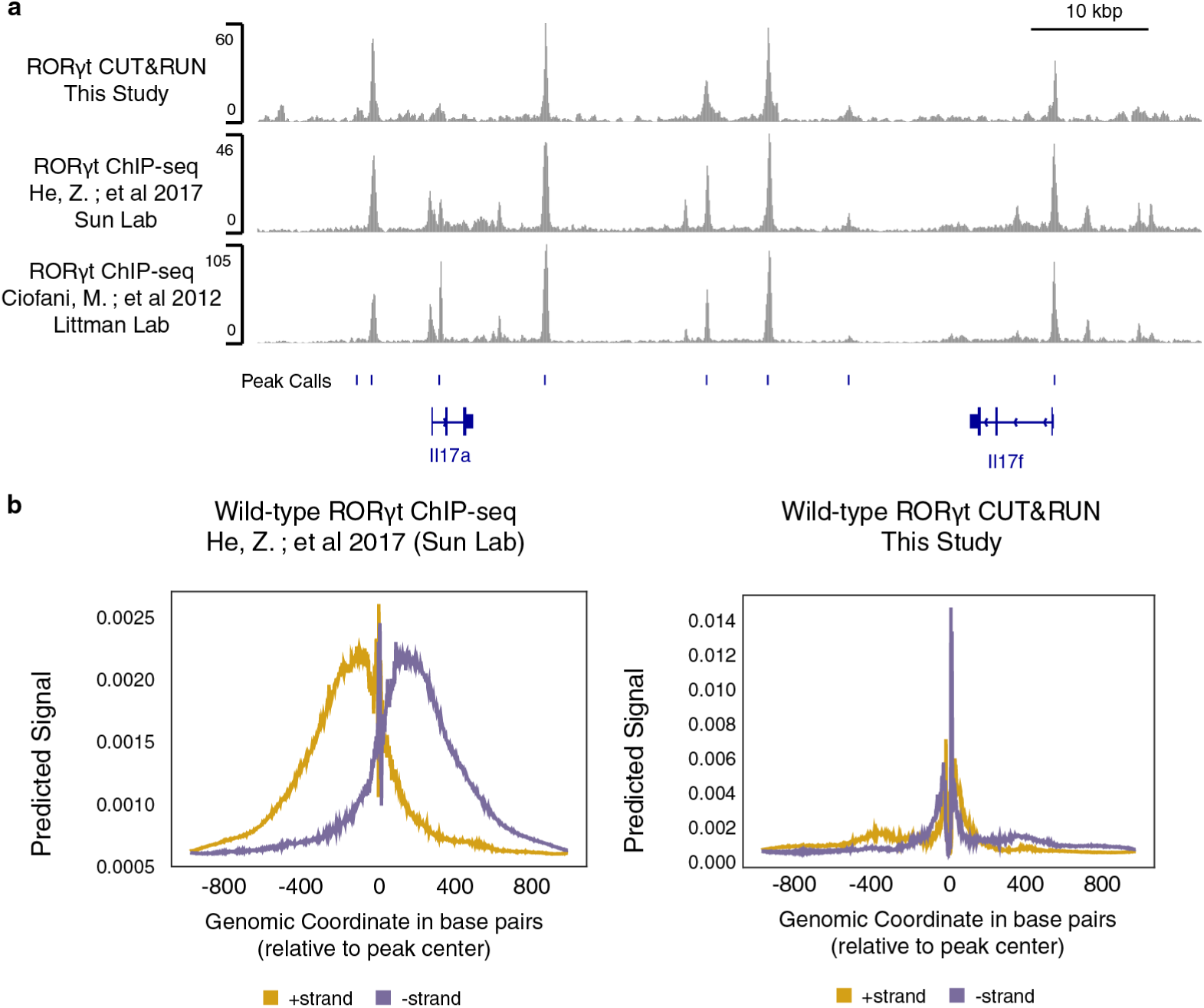
BPnet model analysis and novel motif discovery. (**a**) RORγt genomic occupancy from our CUT&RUN analysis is compared to previously published datasets. BPNet models trained on previously published ChIP-seq datasets and our CUT&RUN datasets. (**b**) BPNet models were tested for sensitivity for a cRORE using marginalization.

**Extended Data Table 1:** HDX-MS data collection summary statistics. Full data export is available through PRIDE entry PXD079146.

| Data Set | RORyt ± Nucleosome |
| --- | --- |
| HDX reaction details | 15 mM HEPES pH 7.9, 100 mM NaCl, 1 mM DTT |
| HDX time course (s) | 10, 30, 60, 300, 900, 3600 |
| HDX control samples | T0 (undeuterated) control |
| T infinity (maximally deuterated) Calculation | estimated from input recovery estimate of 0.7 and Deuterium solution concentration of 0.8 |
| **RORyt** |  |
| # of Peptides | 35 |
| Sequence coverage | 52.5% |
| Average peptide length / Redundancy | 1 / 2.966 |
| Replicates (biological or technical) | 3 (technical) |
| Repeatability | 6.34 % (average standard deviation) |

| Data Set | Nucleosome ± RORyt |
| --- | --- |
| HDX reaction details | 15 mM HEPES pH 7.9, 100 mM NaCl, 1 mM DTT |
| HDX time course (s) | 10, 30, 60, 300, 900, 3600 |
| HDX control samples | T0 (undeuterated) control |
| T infinity (maximally deuterated) Calculation | estimated from input recovery estimate of 0.7 and Deuterium solution concentration of 0.8 |
| **Histone H2A** |  |
| # of Peptides | 35 |
| Sequence coverage | 90.7% |
| Average peptide length / Redundancy | 7 / 5.291 |
| Replicates (biological or technical) | 3 (technical) |
| Repeatability | 4.93 % (average standard deviation) |
| **Histone H2B** |  |
| # of Peptides | 47 |
| Sequence coverage | 84.8% |
| Average peptide length / Redundancy | 7 / 9.085 |
| Replicates (biological or technical) | 3 (technical) |
| Repeatability | 2.84 % (average standard deviation) |
| **Histone H3** |  |
| # of Peptides | 42 |
| Sequence coverage | 80.0% |
| Average peptide length / Redundancy | 3 / 5.417 |
| Replicates (biological or technical) | 3 (technical) |
| Repeatability | 4.08 % (average standard deviation) |
| **Histone H4** |  |
| # of Peptides | 29 |
| Sequence coverage | 100.0% |
| Average peptide length / Redundancy | 5 / 4.882 |
| Replicates (biological or technical) | 3 (technical) |
| Repeatability | 3.20 % (average standard deviation) |

**Extended Data Table 2:** CryoEM data collection, processing, and modeling statistics.

| Name |  | RORyt:Nucleosome |  | Nucleosome |  |
| --- | --- | --- | --- | --- | --- |
| Data collection |  |  |  |  |  |
| Microscope |  | Titan Krios (FEI) |  | Titan Krios G3i (Thermo) |  |
| Detector |  | K3 (Gatan) |  | Falcon 4i (Thermo) |  |
| Magnification |  | 29,000X |  | 147,000X |  |
| Voltage (source) |  | 300 keV (FEG) |  | 300 keV (cold FEG) |  |
| Defocus range |  | `-0.7 to -2.5 μm |  | `-0.7 to -2.5 μm |  |
| Total Dose |  | 49.23 e<sup>-</sup>/Å<sup>2</sup> |  | 50.0 e<sup>-</sup>/Å<sup>2</sup> |  |
| Dose rate |  | 0.7 e<sup>-</sup>/pixel/s |  | 0.83 e<sup>-</sup>/pixel/s |  |
| Pixel size |  | 0.53 Å |  | 0.954 Å |  |
| Number of frames |  | 72 |  | 60 |  |
| Automation Software |  | serialEM |  | EPU |  |
| Movies collected |  | 9668 |  | 4058 |  |
| Data Processing |  |  |  |  |  |
| Data processing software |  | CryoSPARC v4.6.2 /<br>Relion v5.0b |  | CryoSPARC v4.6.2 /<br>Relion v5.0b |  |
| Total Extracted Particles |  | 1408179 |  | 1138327 |  |
| Final Particle Stack |  | 300923 |  | 247702 |  |
| Gold standard FSC |  | 2.61 Å |  | 3.05 Å |  |
| 3D FSC range |  | 2.74-2.43 Å |  | 3.34-2.79 Å |  |
| Sampling Compensation Factor |  | 0.882 |  | 0.728 |  |
| EMDB Ascension |  | EMD-77866 |  | EMD-77867 |  |
| Modeling |  |  |  |  |  |
| Software |  | Alphafold3 / ISOLDE<br>v1.8 / ChimeraX v 1.8 /<br>Phenix 1.21 |  | Alphafold3 / ISOLDE<br>v1.8 / ChimeraX v 1.8 /<br>Phenix 1.21 |  |
| PDB ID |  | pdb_000036UG |  | pdb_000036UH |  |
| Map-to-model FSC (cut off 0.5) |  | 2.90 Å |  | 3.31 Å |  |
| All-Atom | Clashscore, all atoms: | 5.48 |  | 7.76 |  |
| Protein | Poor rotamers | 0 | 0.00% | 7 | 1.05% |
| Geometry | Favored rotamers | 568 | 83.53% | 559 | 83.93% |
|  | Ramachandran outliers | 1 | 0.13% | 0 | 0.00% |
|  | Ramachandran favored | 785 | 98.25% | 782 | 98.86% |
|  | Rama distribution Z-score | 1.51 ± 0.28 |  | 1.43 ± 0.27 |  |
|  | MolProbity score^ | 1.3 |  | 1.44 |  |
|  | Cβ deviations >0.25Å | 0 | 0.00% | 0 | 0.00% |
|  | Bad bonds: | 3 / 6539 | 0.05% | 0 / 6489 | 0.00% |
|  | Bad angles: | 7 / 8783 | 0.08% | 0 / 8698 | 0.00% |
| Peptide Omegas | Cis Prolines: | 0 / 22 | 0.00% | 0 / 23 | 0.00% |
| Nucleic Acid | Bad bonds: | 0 / 6438 | 0.00% | 0 / 6668 | 0.00% |
| Geometry | Bad angles: | 0 / 9932 | 0.00% | 0 / 10287 | 0.00% |
| Low-resolution Criteria | CaBLAM outliers | 10 | 1.30% | 5 | 0.60% |
|  | CA Geometry outliers | 2 | 0.26% | 1 | 0.13% |

**Extended Data Table 3:**
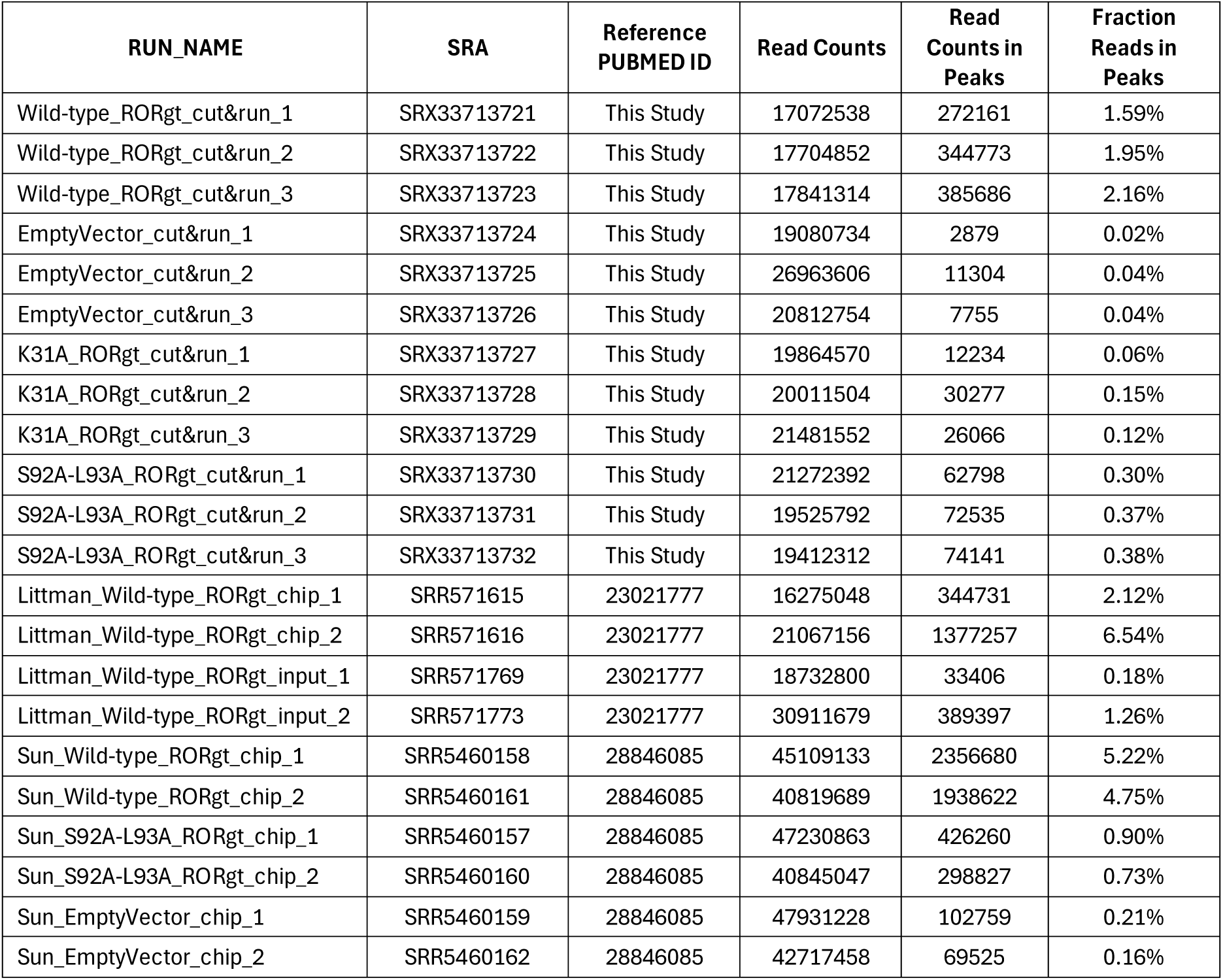
CUT&RUN and ChIP-seq datasets analyzed in this study.

| **RUN_NAME** | **SRA** | **Reference**<br>**PUBMED ID** | **Read Counts** | **Read**<br>**Counts in**<br>**Peaks** | **Fraction**<br>**Reads in**<br>**Peaks** |
| --- | --- | --- | --- | --- | --- |
| Wild-type_RORgt_cut&run_1 | SRX33713721 | This Study | 17072538 | 272161 | 1.59% |
| Wild-type_RORgt_cut&run_2 | SRX33713722 | This Study | 17704852 | 344773 | 1.95% |
| Wild-type_RORgt_cut&run_3 | SRX33713723 | This Study | 17841314 | 385686 | 2.16% |
| EmptyVector_cut&run_1 | SRX33713724 | This Study | 19080734 | 2879 | 0.02% |
| EmptyVector_cut&run_2 | SRX33713725 | This Study | 26963606 | 11304 | 0.04% |
| EmptyVector_cut&run_3 | SRX33713726 | This Study | 20812754 | 7755 | 0.04% |
| K31A_RORgt_cut&run_1 | SRX33713727 | This Study | 19864570 | 12234 | 0.06% |
| K31A_RORgt_cut&run_2 | SRX33713728 | This Study | 20011504 | 30277 | 0.15% |
| K31A_RORgt_cut&run_3 | SRX33713729 | This Study | 21481552 | 26066 | 0.12% |
| S92A-L93A_RORgt_cut&run_1 | SRX33713730 | This Study | 21272392 | 62798 | 0.30% |
| S92A-L93A_RORgt_cut&run_2 | SRX33713731 | This Study | 19525792 | 72535 | 0.37% |
| S92A-L93A_RORgt_cut&run_3 | SRX33713732 | This Study | 19412312 | 74141 | 0.38% |
| Littman_Wild-type_RORgt_chip_1 | SRR571615 | 23021777 | 16275048 | 344731 | 2.12% |
| Littman_Wild-type_RORgt_chip_2 | SRR571616 | 23021777 | 21067156 | 1377257 | 6.54% |
| Littman_Wild-type_RORgt_input_1 | SRR571769 | 23021777 | 18732800 | 33406 | 0.18% |
| Littman_Wild-type_RORgt_input_2 | SRR571773 | 23021777 | 30911679 | 389397 | 1.26% |
| Sun_Wild-type_RORgt_chip_1 | SRR5460158 | 28846085 | 45109133 | 2356680 | 5.22% |
| Sun_Wild-type_RORgt_chip_2 | SRR5460161 | 28846085 | 40819689 | 1938622 | 4.75% |
| Sun_S92A-L93A_RORgt_chip_1 | SRR5460157 | 28846085 | 47230863 | 426260 | 0.90% |
| Sun_S92A-L93A_RORgt_chip_2 | SRR5460160 | 28846085 | 40845047 | 298827 | 0.73% |
| Sun_EmptyVector_chip_1 | SRR5460159 | 28846085 | 47931228 | 102759 | 0.21% |
| Sun_EmptyVector_chip_2 | SRR5460162 | 28846085 | 42717458 | 69525 | 0.16% |

